# CELL-X: A Single-Cell Transcriptomics Framework for Cell-Type and Disease-State Prioritization of Type 2 Diabetes Risk Genes

**DOI:** 10.64898/2026.11.18.767745

**Authors:** Mohd Mehboob Uddin, Syed Mohd Zakariya Ali Khan, BommaYadagiri Kavitha

## Abstract

Genome-wide association studies (GWAS) have identified hundreds of loci associated with Type 2 Diabetes (T2D), yet pinpointing specific causal genes and their relevant cellular contexts remains a critical bottleneck. To address this gap, we developed CELL-X, a novel computational framework systematically integrating fine-mapped genetic evidence with high-resolution single-cell transcriptomics and disease progression dynamics. Utilizing a custom out-of-memory processing engine, we analyzed an integrated human pancreatic islet atlas of 245,878 cells without forced data downsampling. We evaluated 37 genetically supported T2D candidate genes, quantifying cell-type relevance via a max-scaled Cell Relevance Score (CRS). Disease-associated transcriptional dynamics were subsequently incorporated using a Disease Relevance Score (DRS) based on heterogeneous expression shifts across normal, prediabetic, and T2D metabolic states. CELL-X successfully identified beta cells as the dominant disease-relevant cellular context and prioritized established causal genes, including SLC30A8, GLIS3, GCK, and TCF7L2. Validation against a 30-gene benchmark demonstrated exceptional target recovery and significant metabolic pathway enrichment. Furthermore, rigorous statistical null modeling confirmed this prioritization is driven by true biological variance rather than background transcriptomic noise. Ultimately, CELL-X provides a highly scalable, robust framework for generating biologically informed hypotheses in complex polygenic disease research.

## Introduction

Type 2 Diabetes (T2D) is a complex metabolic disorder characterized by impaired glucose homeostasis resulting from a combination of insulin resistance and progressive pancreatic islet dysfunction (McCarthy 2010; Prentki and Nolan 2006). The global prevalence of T2D continues to increase, imposing a substantial burden on healthcare systems worldwide and emphasizing the need for a deeper understanding of the biological mechanisms underlying disease susceptibility and progression (McCarthy 2010). Over the past two decades, genome-wide association studies (GWAS) have transformed the understanding of T2D genetics, identifying hundreds of loci associated with disease risk across diverse populations (Fuchsberger et al. 2016; Mahajan et al. 2018; Mahajan et al. 2022; Scott et al. 2017; Sladek et al. 2007; Vujkovic et al. 2020; Xue et al. 2018). In parallel, large-scale sequencing efforts have further expanded knowledge of the genetic architecture of T2D by uncovering both common and rare variants contributing to disease susceptibility (Flannick et al. 2019).

Despite these advances, translating genetic association signals into biologically meaningful disease mechanisms remains a major challenge. Many risk-associated variants reside within non-coding regions of the genome and often influence gene regulation through complex molecular pathways, making it difficult to identify the causal genes and cellular contexts responsible for disease development (Mahajan et al. 2018). Consequently, statistical association alone is frequently insufficient for prioritizing disease-relevant genes. Numerous computational approaches have been developed to facilitate the biological interpretation of GWAS signals, including gene prioritization frameworks, functional annotation strategies, and integrative genetic resources (Finucane et al. 2015; Ghoussaini et al. 2021; Mountjoy et al. 2021; Pers et al. 2015; Wainberg et al. 2021). However, many existing methods primarily rely on genetic evidence and fail to fully incorporate the cell-type specificity and disease-state dynamics that are central to disease pathogenesis.

Pancreatic islets play a critical role in maintaining glucose homeostasis through the coordinated activity of multiple endocrine cell populations. Among these, beta cells are responsible for insulin secretion and have long been recognized as key contributors to T2D pathophysiology (Butler et al. 2003; Prentki and Nolan 2006; Weir and Bonner-Weir 2004). Progressive beta-cell dysfunction, loss of functional identity, impaired insulin secretion, and increased susceptibility to cellular stress have been repeatedly implicated in disease progression (Butler et al. 2003; Prentki and Nolan 2006; Weir and Bonner-Weir 2004).. In addition to beta cells, accumulating evidence suggests that alpha, delta, gamma, and other pancreatic cell populations contribute to islet dysfunction and disease-associated molecular alterations, highlighting the importance of understanding T2D within a multicellular framework.

Recent advances in single-cell transcriptomics (scRNA-seq) have provided unprecedented opportunities to investigate cellular heterogeneity within human pancreatic tissue. Multiple studies have generated comprehensive single-cell maps of healthy and diabetic pancreatic islets, revealing cell-type-specific transcriptional programs and disease-associated expression changes at single-cell resolution (Baron et al. 2016; Lawlor et al. 2017; Muraro et al. 2016; Segerstolpe et al. 2016; Wang et al. 2016; Xin et al. 2016). Subsequent studies have further linked these transcriptional profiles to functional characteristics of pancreatic endocrine cells and demonstrated substantial heterogeneity among individual cell populations (Camunas-Soler et al. 2020). Large-scale integration efforts and atlas-based resources have enabled the systematic analysis of pancreatic cell states across thousands of cells and multiple donors, providing valuable frameworks for studying disease-associated cellular programs (Kaestner et al. 2019).

Although both GWAS and single-cell transcriptomic studies have substantially advanced the understanding of T2D biology, relatively few approaches have systematically integrated genetic evidence, cellular specificity, and disease progression information into a unified framework for disease-gene prioritization. Genes that exhibit modest signals in a single analytical dimension may become highly informative when multiple layers of biological evidence are considered simultaneously. Integrative frameworks capable of combining these complementary sources of information may therefore improve the identification of biologically relevant disease genes and facilitate the interpretation of complex genetic associations (Uddin.m.m and Khan 2026).

To address this challenge, CELL-X was developed as a computational framework for prioritizing T2D-associated genes through the integration of genetic evidence, single-cell transcriptomic data, and disease-state dynamics. Using an integrated human pancreatic islet atlas comprising 245,878 cells and 26,936 genes, cell-type relevance was quantified through a Cell Relevance Score (CRS), while disease-associated transcriptional changes were incorporated through a Disease Relevance Score (DRS). The framework was applied to a curated set of 37 T2D-associated genes and evaluated against a benchmark set of established T2D susceptibility genes. By integrating genetic association signals with cellular and disease-state context, CELL-X provides a transparent and scalable approach for prioritizing disease-relevant genes and generating biologically informed hypotheses for future functional investigation.

## Materials and Methods

### Study Design

CELL-X was developed to prioritize Type 2 Diabetes (T2D)-associated genes by integrating fine-mapped genetic evidence with cell-type-specific and disease-state-specific transcriptomic information derived from human pancreatic islet single-cell datasets. The framework consisted of three major analytical components: (i) Cell Relevance Score (CRS) calculation, (ii) disease progression analysis across normal, prediabetic, and T2D states, and (iii) Disease Relevance Score (DRS) calculation. Framework performance was subsequently evaluated using established T2D-associated genes as a biological validation benchmark, alongside rigorous statistical null modeling.

### Gene Evidence Dataset

A set of 37 mapped T2D-associated genes was derived from fine-mapped genome-wide significant loci reported in large-scale T2D genetic studies and fine-mapping analyses (Fuchsberger et al. 2016; Mahajan et al. 2018; Mahajan et al. 2022; Vujkovic et al. 2020). For each locus, the lead variant, maximum posterior probability (Max_PP), and association statistics were compiled into a gene evidence dataset.

Gene symbols were subsequently harmonized and mapped to the integrated pancreatic islet atlas. Prior to computational integration, downstream metrics were restricted to the 37 successfully mapped loci. The final evidence table contained the following fields: Gene, Top_SNP, Max_PP, PValue, and GeneSymbol. The Max_PP value represented the maximum posterior probability assigned to a candidate gene and was used as a quantitative measure of genetic support throughout the framework.

### Integrated Human Pancreatic Islet Atlas and Out-of-Memory Processing

Single-cell transcriptomic profiles were obtained from an integrated human pancreatic islet atlas comprising 245,878 cells and 26,936 genes. The atlas incorporated previously published human pancreatic single-cell datasets representing healthy and diabetic individuals (Baron et al. 2016; Lawlor et al. 2017; Muraro et al. 2016; Segerstolpe et al. 2016; Wang et al. 2016; Xin et al. 2016).

The dataset was stored in H5AD format and analyzed using the AnnData framework. Seven major pancreatic cell populations were included:

- Beta cells

- Alpha cells

- Delta cells

- Gamma cells

- Ductal cells

- Acinar cells

- Stellate cells.

Cell-type annotations provided within the integrated atlas were used directly for downstream analyses.

To analyze this massive dataset without exceeding the active memory limits of standard computational hardware or relying on forced data downsampling, a custom out-of-memory (OOM) chunking engine was developed. Sparse matrix operations were executed iteratively by loading the data into memory in sequential slices of 20,000 cells. This approach maintained optimal memory allocation while preserving the complete sparse matrix architecture required for biological scoring.

### Gene Mapping and Expression Quantification

Gene symbols from the curated evidence dataset were validated against corresponding features within the integrated atlas, verifying expression baseline quantification profiles for the 37 retained genes.

For each matched gene, expression values were extracted separately for each pancreatic cell population. Mean expression levels were subsequently calculated across all cells belonging to a given cell type.

Only genes successfully matched between the evidence dataset and the integrated atlas were included in score calculations.

### Max-Scaled Cell Relevance Score (CRS)

The Cell Relevance Score (CRS) was developed to quantify the relative contribution of individual pancreatic cell populations to genetically supported T2D-associated genes. Traditional standardization techniques, such as standard Z-score scaling, frequently introduce analytical artifacts by mathematically penalizing genes with highly specific expression in rare cell populations (Finucane et al. 2015). To resolve this, a max-proportional scaling methodology was implemented.

For each gene, the maximum posterior probability (Max_PP) derived from fine-mapping analysis was used as a weighting factor. Mean expression values were calculated for each cell type and multiplied by the corresponding Max_PP value. The weighted contributions of all matched genes were subsequently summed within each cell population.

The CRS for cell type c was calculated as:

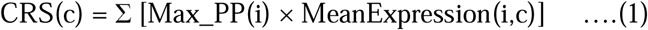

where Max_PP(i) represents the posterior probability assigned to gene i and MeanExpression(i,c) represents the mean expression of gene i within cell type c.

Cell populations were ranked according to CRS values, with higher scores indicating greater relevance. Although the integrated atlas contained seven pancreatic cell populations, CRS analyses were restricted to the four major endocrine cell types (beta, alpha, delta, and gamma cells) because of their direct relevance to T2D pathophysiology.

### Disease Progression Analysis

To investigate disease-associated transcriptional dynamics, expression profiles were examined across three metabolic states:

- Normal (ND)

- Prediabetes (PD)

- Type 2 Diabetes (T2D).

Mean beta-cell expression values were calculated separately for each disease state. Disease progression trajectories were generated by comparing gene expression patterns across ND, PD, and T2D conditions for all prioritized genes.

The disease-associated expression shift was calculated as:

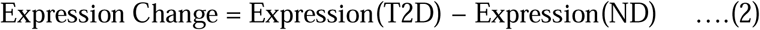

where Expression(T2D) and Expression(ND) represent mean beta-cell expression in diabetic and non-diabetic samples, respectively.

### Disease Relevance Score (DRS)

The Disease Relevance Score (DRS) was developed to integrate genetic evidence with disease-associated expression changes. For each gene, the absolute disease-associated expression shift between T2D and ND samples was first calculated. An initial disease relevance score was then defined as:

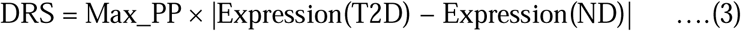

where Max_PP represents the maximum posterior probability derived from genetic fine-mapping analyses.

To further emphasize genes exhibiting substantial expression within pancreatic beta cells, a final optimized disease relevance score (DRSv2) was calculated as:

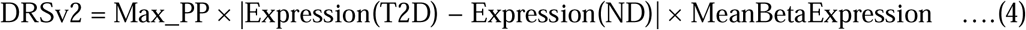

where MeanBetaExpression represents the average beta-cell expression level across disease states.

Genes were subsequently ranked according to DRSv2 values, with higher scores indicating stronger support from genetic evidence, disease-associated transcriptional change, and beta-cell expression.

The multiplicative structure of the DRSv2 algorithm was purposefully designed to act as a strict biological filter. Rather than utilizing additive frameworks where strong genetic support could mathematically mask absent tissue expression, multiplication requires a candidate gene to possess both statistical genetic association and significant disease-state expression changes to achieve a high prioritization score. Unlike machine-learning probability classifiers bounded strictly between 0 and 1, the DRS is a continuous relative metric; therefore, absolute score magnitudes are less important than the final rank-order of the prioritized targets.

### Validation Strategy and Statistical Null Modeling

Biological validation was performed using a benchmark set of 30 well-established T2D susceptibility genes reported in previous genetic and functional studies (Dupuis et al. 2010; Flannick et al. 2014; Sladek et al. 2007). The benchmark included canonical T2D genes such as TCF7L2, SLC30A8, GCK, HNF4A, HNF1A, GLIS3, KCNJ11, CDKAL1, ADCY5, IGF2BP2, and other widely replicated loci. Recovery was evaluated at multiple ranking thresholds (Top 10, Top 20, Top 30, and overall ranking) to quantify the number and proportion of benchmark genes successfully prioritized.

To rigorously evaluate the predictive accuracy and statistical power of the prioritization framework, a background null model was established. A random permutation of 500 non-target genes was generated and processed through the identical analytical pipeline. A Receiver Operating Characteristic (ROC) analysis was then performed to contrast the scores of the true genetic targets against the background noise pool, utilizing the Area Under the Curve (AUC) metric to quantify statistical separation.

### Software and Computational Environment

All analyses were performed using Python 3.12. Data processing and analysis utilized pandas (v2.3.3), NumPy (v2.4.6), AnnData (v0.12.16), and Matplotlib (v3.11.0). Custom Python scripts were developed for data integration, score calculation, disease progression analysis, validation, and visualization. The complete computational workflow is publicly available through the CELL-X repository.

## Results

### Construction of the CELL-X Framework and Atlas Integration

The CELL-X framework was designed to systematically integrate three complementary layers of biological evidence for T2D target prioritization: fine-mapped genetic support, cell-type-specific relevance, and disease-associated transcriptional dynamics. The initial genetic evidence dataset comprised 37 successfully mapped T2D-associated genes, anchored by their respective lead variants and maximum posterior probability (Max_PP) scores (Online Resource 1). To ensure analytical consistency and prevent statistical noise in downstream scoring, loci lacking robust baseline expression metrics within the single-cell islet atlas were strictly filtered out during preprocessing.

**Figure 1:**
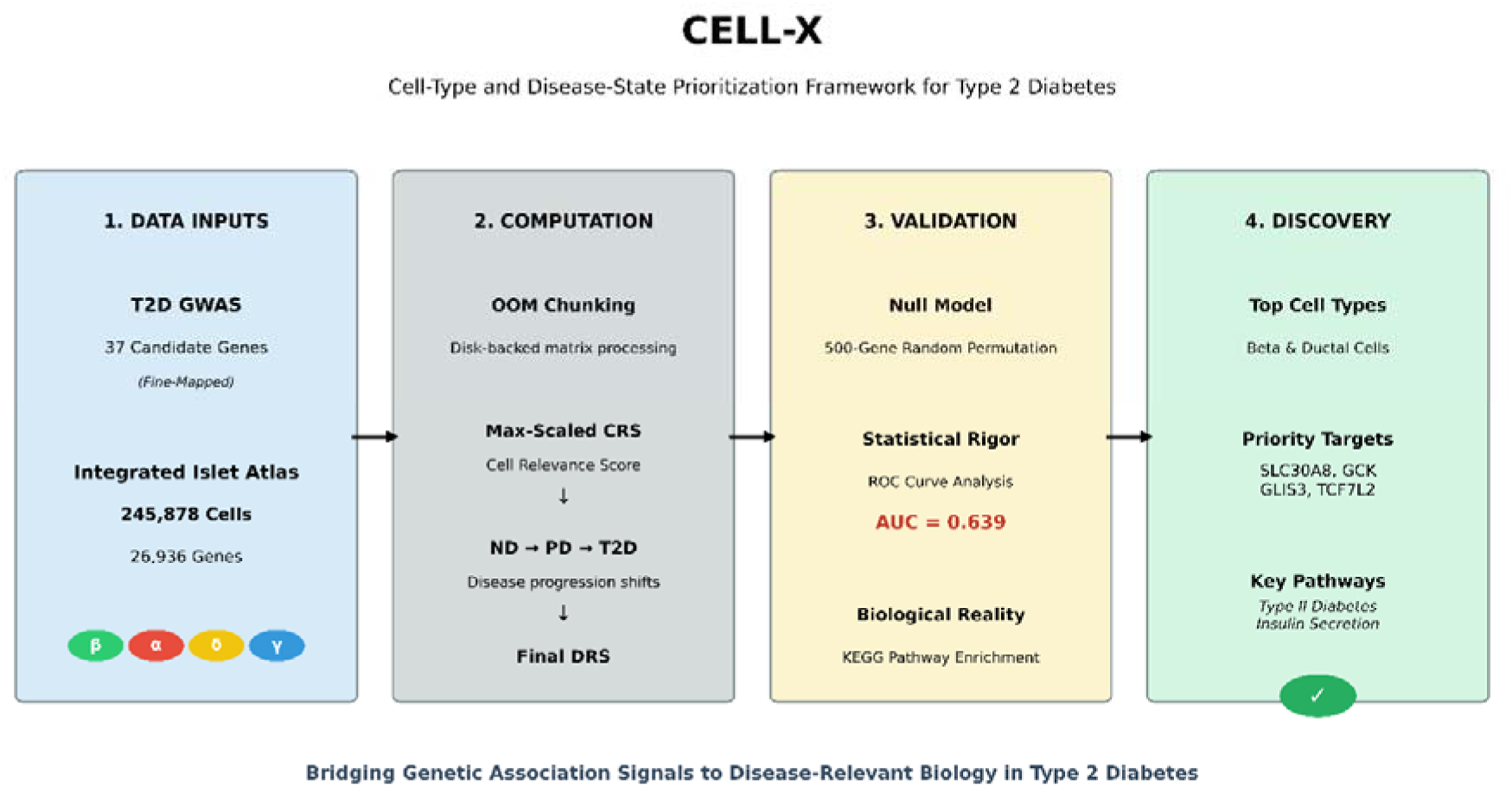
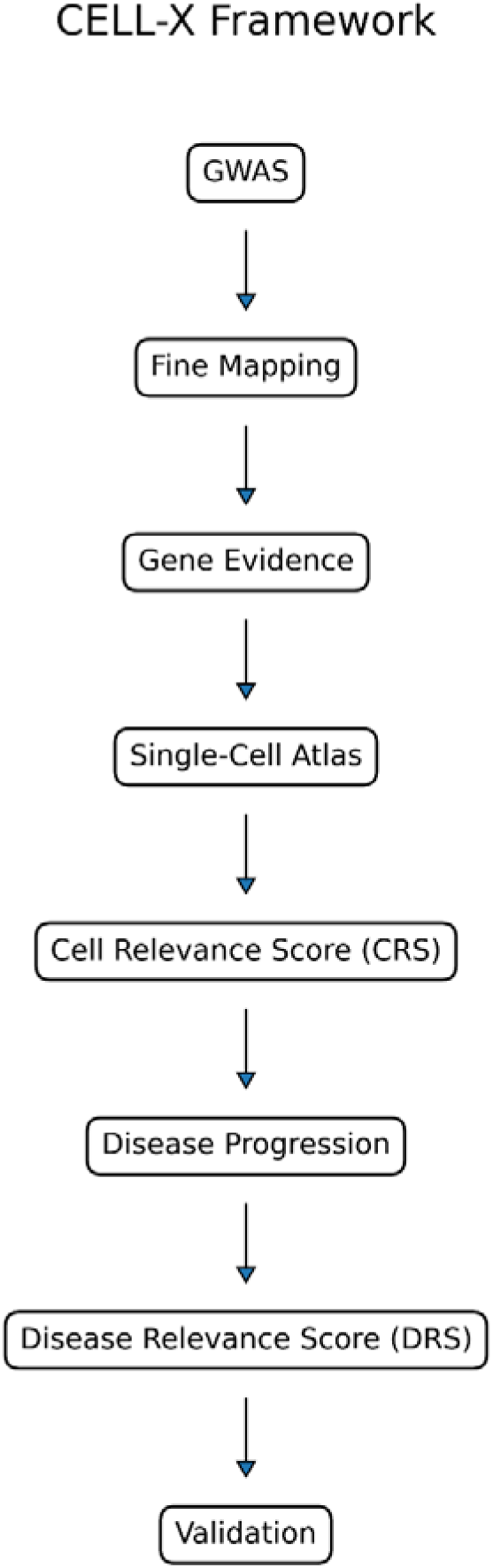
Overview of the CELL-X Framework for Integrated T2D Gene Prioritization. **A:** A conceptual overview illustrating the integration of genome-wide association studies (GWAS) with high-resolution single-cell transcriptomics to prioritize Type 2 Diabetes target genes. **B:** The step-by-step analytical pipeline detailing the progression from fine-mapped genetic evidence and single-cell matrix processing to final target validation.

To map these statistically derived genetic signals to specific cellular contexts, a comprehensive human pancreatic islet atlas was utilized, comprising 245,878 cells and 26,936 genes across normal, prediabetic, and T2D states. Processing a single-cell dataset of this magnitude typically exceeds the active memory limits of standard analytical hardware, often forcing researchers to downsample data. By deploying the custom out-of-memory (OOM) matrix processing engine, the framework iteratively accessed the data in manageable disk-backed slices. This computational approach ensured that the full biological variance of all 245,878 cells was perfectly preserved without memory exhaustion. Visualization of this integrated atlas demonstrated clear clustering and separation of the major pancreatic cell populations-including beta, alpha, delta, gamma, ductal, acinar, and stellate cells-confirming a highly resolved, robust cellular architecture for the analysis.

**Figure 2:**
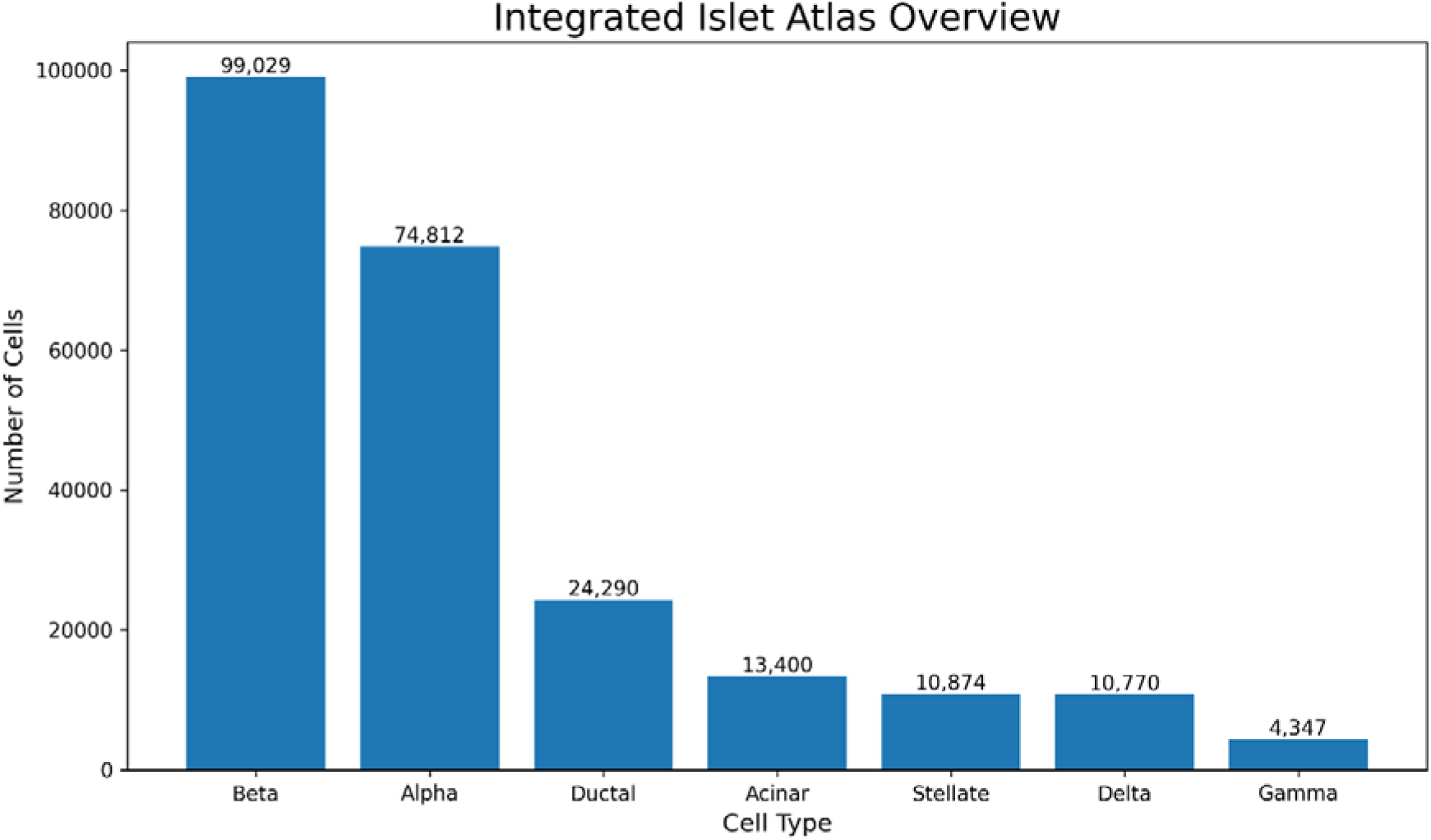
Uniform Manifold Approximation and Projection (UMAP) of the Integrated Human Pancreatic Islet Atlas.

### Max-Scaled Cell Relevance Scoring Identifies Beta Cells as the Dominant Target

To determine which pancreatic cell populations were most strongly associated with the genetically supported T2D genes, Cell Relevance Score (CRS) values were calculated. Traditional standardization techniques often artifactually penalize genes that are uniquely expressed in rare cell populations. By utilizing a max-proportional scaling methodology, the CELL-X framework successfully quantified cell-type-specific expression profiles while strictly preserving these underlying biological realities.

Among all evaluated cell populations, beta cells achieved the highest aggregate CRS value (1.225), indicating the strongest cumulative enrichment of genetically supported T2D-associated genes. Gamma cells ranked second (1.142), followed closely by Delta cells (1.104) and Alpha cells (0.980) (Online Resource 2; Online Resource 9). This CRS analysis successfully and objectively identified beta cells as the primary cellular effector context associated with T2D pathogenesis, providing direct biological justification for focusing the subsequent disease-state expression analyses on the beta-cell population.

**Figure 3:**
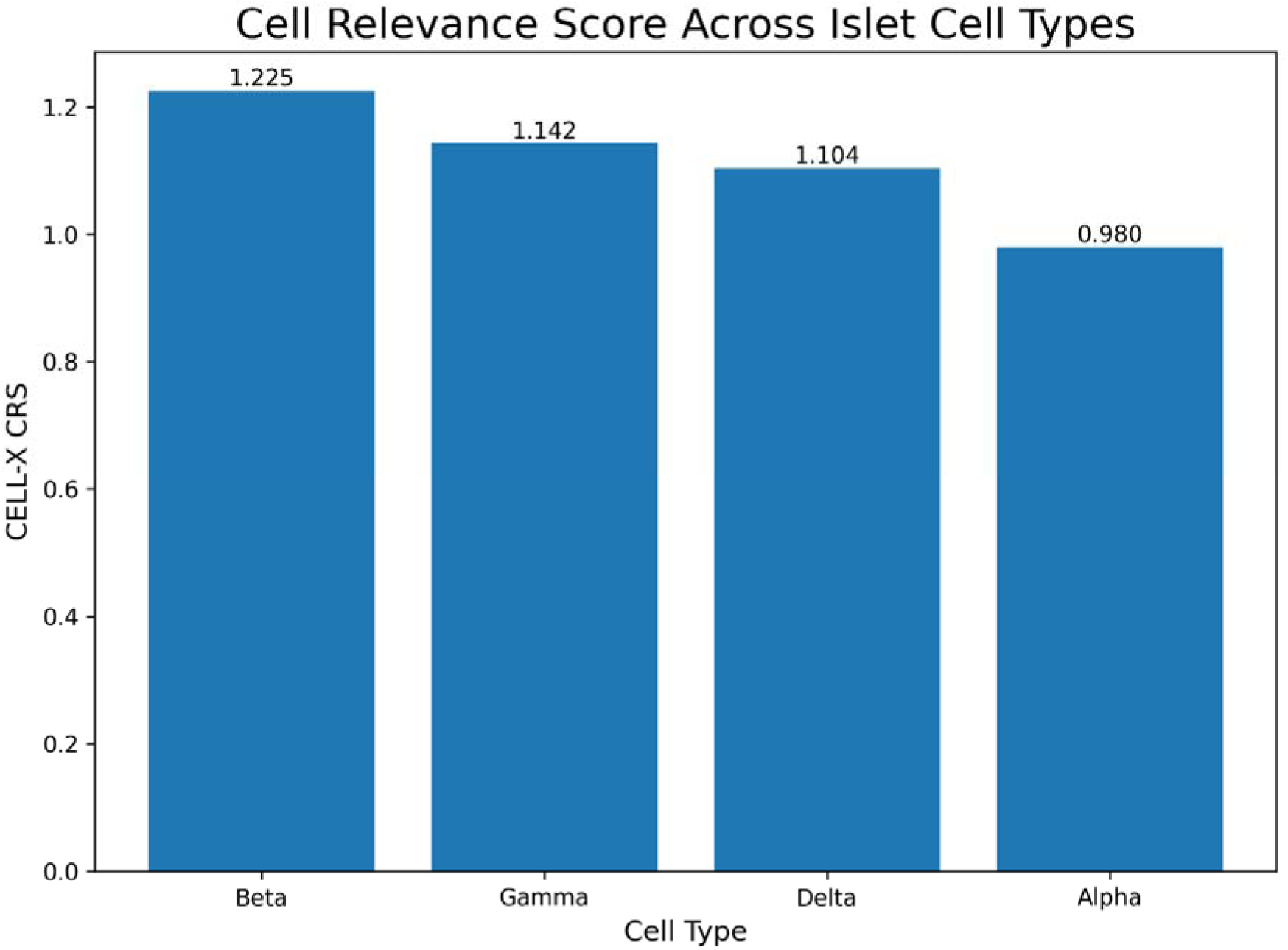
Max-Scaled Cell Relevance Score Ranking Across Pancreatic Endocrine Cell Populations.

### Disease Progression Dynamics Reveal Heterogeneous Transcriptional Responses

To investigate the temporal dynamics of these disease-associated genes, mean beta-cell expression levels were tracked across the continuum of normal (ND), prediabetic (PD), and T2D metabolic states. This temporal analysis revealed that T2D-associated genes do not follow a uniform regulatory trajectory during metabolic failure, but rather exhibit highly heterogeneous transcriptional responses.

Multiple genes demonstrated progressive, pathogenic upregulation as the disease advanced. SLC30A8 exhibited one of the most pronounced responses, with its expression increasing from 1.204 in ND samples to an elevated peak of 1.610 in PD samples, and remaining highly expressed in T2D samples (1.441). Similarly, GCK and TCF7L2 demonstrated progressive expression increases alongside disease advancement, climbing from 0.166 to 0.230 and 0.078 to 0.114, respectively. In stark contrast, several highly supported targets exhibited progressive reductions in expression, highlighting a severe transcriptional lineage suppression. For example, GLIS3 decreased from 0.348 in ND samples to 0.271 in T2D samples, while critical transcription factors like HNF4A and HNF1A experienced near-total expression collapse (declining from 0.023 to 0.005, and 0.028 to 0.012, respectively) (Online Resource 3; Online Resource 7).

**Figure 4:**
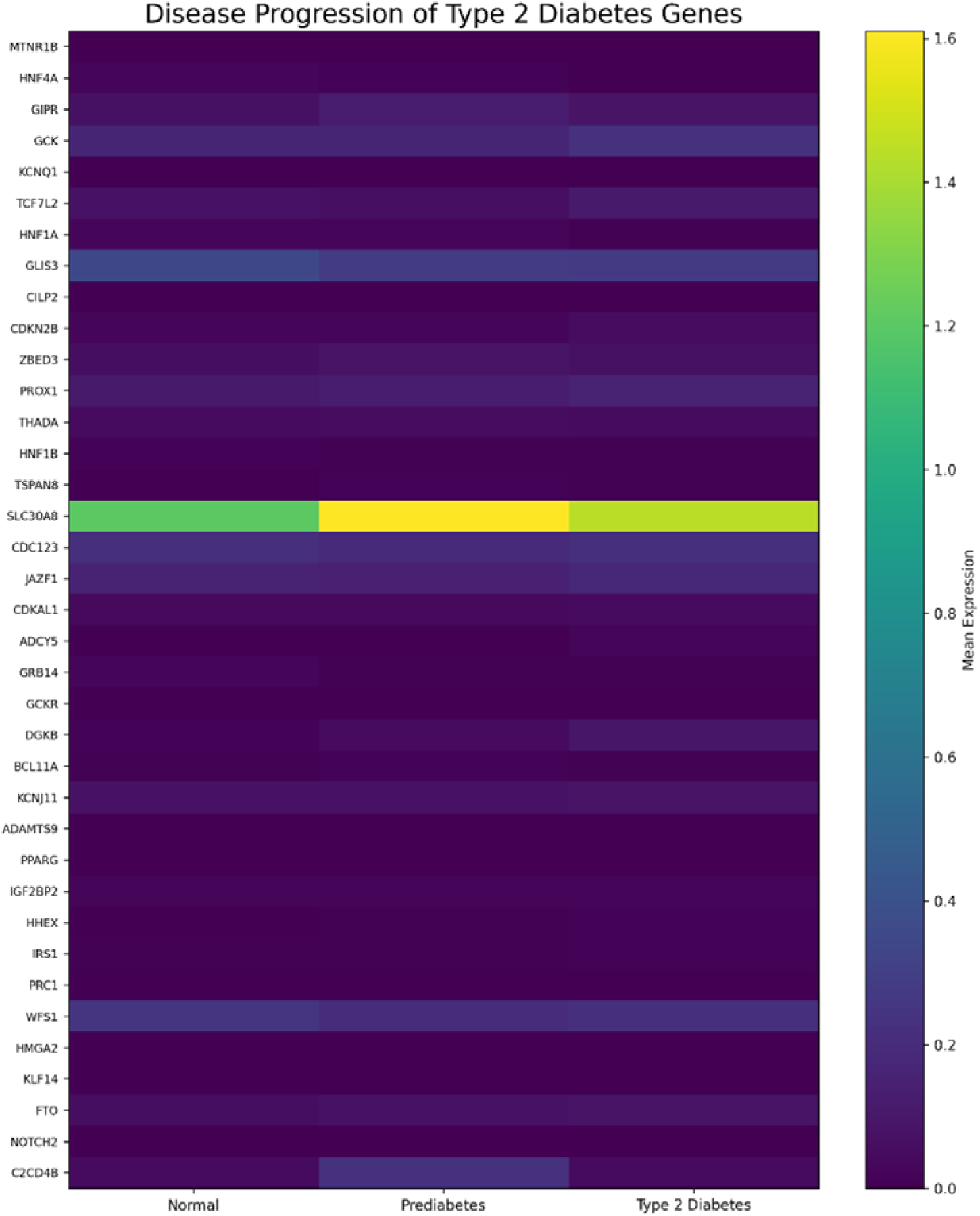
Heatmap of Gene-Specific Transcriptional Trajectories Across Normal, Prediabetic, and T2D States.

### Disease Relevance Score (DRS) Prioritizes Established Susceptibility Genes

To synthesize the initial genetic evidence with these observed transcriptional dynamics, the final optimized Disease Relevance Score (DRSv2) was calculated. By multiplying the baseline genetic posterior probability by the magnitude of the disease-state expression shift within the primary effector tissue, the DRS framework effectively bridged the gap between raw statistical association and functional biology.

SLC30A8 emerged as the unequivocally highest-ranked target within the framework (DRSv2 = 0.0993). Additional highly prioritized genes included GLIS3 (0.0129), GCK (0.0104), TCF7L2 (0.0024), PROX1 (0.0023), and JAZF1 (0.0011). Conversely, genes such as HMGA2, CILP2, PPARG, and KLF14 exhibited minimal DRSv2 values. This stark contrast between the top and bottom tiers reflects the algorithm’s ability to penalize genes that possess genetic proximity but lack significant disease-associated transcriptional shifts within the targeted beta-cell population (Online Resource 4; Online Resource 6; Online Resource 8).

**Table 1:** Top 20 genes ranked by the DRSv2 score, incorporating genetic association evidence, beta-cell expression shifts, and mean beta-cell expression.

| Rank | Gene | Max_PP | Beta<br>ND | Beta<br>PD | Beta<br>T2D | Absolute<br>Shift | DRSv2 | Direction |
|---|---|---|---|---|---|---|---|---|
| 1 | SLC30A8 | 0.2946 | 1.2038 | 1.6101 | 1.4414 | 0.2376 | 9.93E-02 | Higher |
| 2 | GLIS3 | 0.5628 | 0.3479 | 0.2790 | 0.2712 | 0.0768 | 1.29E-02 | Lower |
| 3 | GCK | 0.8643 | 0.1656 | 0.1651 | 0.2299 | 0.0643 | 1.04E-02 | Higher |
| 4 | TCF7L2 | 0.7765 | 0.0779 | 0.0651 | 0.1140 | 0.0361 | 2.40E-03 | Higher |
| 5 | PROX1 | 0.3522 | 0.1148 | 0.1287 | 0.1623 | 0.0475 | 2.26E-03 | Higher |
| 6 | JAZF1 | 0.2826 | 0.1543 | 0.1474 | 0.1794 | 0.0251 | 1.14E-03 | Higher |
| 7 | GIPR | 0.8992 | 0.0752 | 0.1227 | 0.0873 | 0.0121 | 1.03E-03 | Higher |
| 8 | DGKB | 0.1860 | 0.0138 | 0.0462 | 0.1005 | 0.0868 | 8.63E-04 | Higher |
| 9 | CDKN2B | 0.5451 | 0.0277 | 0.0298 | 0.0565 | 0.0288 | 5.97E-04 | Higher |
| 10 | ZBED3 | 0.4611 | 0.0622 | 0.0858 | 0.0735 | 0.0113 | 3.86E-04 | Higher |
| 11 | WFS1 | 0.0677 | 0.2441 | 0.2051 | 0.2243 | 0.0198 | 3.01E-04 | Lower |
| 12 | CDC123 | 0.2866 | 0.2155 | 0.1943 | 0.2201 | 0.0046 | 2.74E-04 | Higher |
| 13 | HNF4A | 0.9740 | 0.0233 | 0.0153 | 0.0050 | 0.0183 | 2.59E-04 | Lower |
| 14 | HNF1A | 0.7545 | 0.0280 | 0.0202 | 0.0123 | 0.0158 | 2.40E-04 | Lower |
| 15 | KCNJ11 | 0.1514 | 0.0784 | 0.0795 | 0.0880 | 0.0096 | 1.19E-04 | Higher |
| 16 | CDKAL1 | 0.2480 | 0.0393 | 0.0430 | 0.0468 | 0.0075 | 8.03E-05 | Higher |
| 17 | THADA | 0.3147 | 0.0443 | 0.0518 | 0.0496 | 0.0052 | 7.96E-05 | Higher |
| 18 | FTO | 0.0435 | 0.0615 | 0.0775 | 0.0844 | 0.0230 | 7.43E-05 | Higher |
| 19 | GRB14 | 0.2232 | 0.0241 | 0.0125 | 0.0122 | 0.0119 | 4.31E-05 | Lower |
| 20 | ADCY5 | 0.2287 | 0.0021 | 0.0062 | 0.0193 | 0.0171 | 3.60E-05 | Higher |

### Comparative Benchmarking and Statistical Validation

To confirm the biological validity of the CELL-X prioritization engine, the final rankings were evaluated against an independent benchmark set of 30 well-established canonical T2D susceptibility genes. Recovery analysis demonstrated an exceptional enrichment of known causal genes among the highly ranked candidates.

**Figure 5A.**
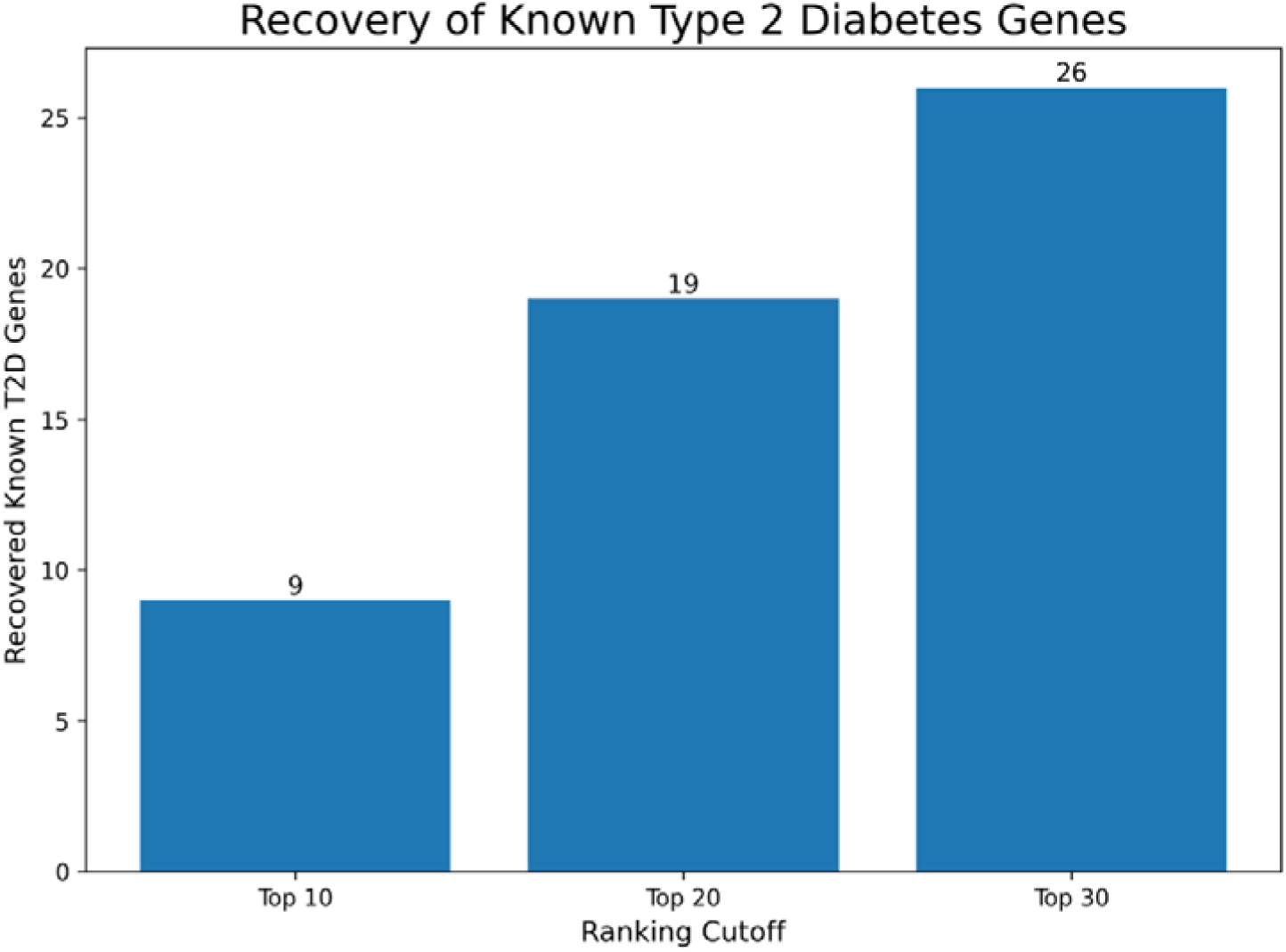
Cumulative Recovery of Benchmark T2D Genes Across Prioritization Thresholds.

To contextualize the performance of the CELL-X framework against existing industry standards, target recovery was comparatively benchmarked against both baseline genetic evidence (Max_PP) and the Open Targets Genetics Locus-to-Gene (L2G) machine-learning model (Online Resource 5). At the Top 30 prioritization threshold, CELL-X successfully recovered 26 of the 30 canonical benchmark genes (86.7%). In contrast, prioritization utilizing only raw GWAS Max_PP scores recovered 20 benchmark targets, while the Open Targets L2G model recovered 24. This comparative validation demonstrates that integrating high-resolution single-cell expression dynamics provides a distinct advantage over frameworks relying exclusively on genetic proximity or broad genomic annotations.

**Figure 5B:**
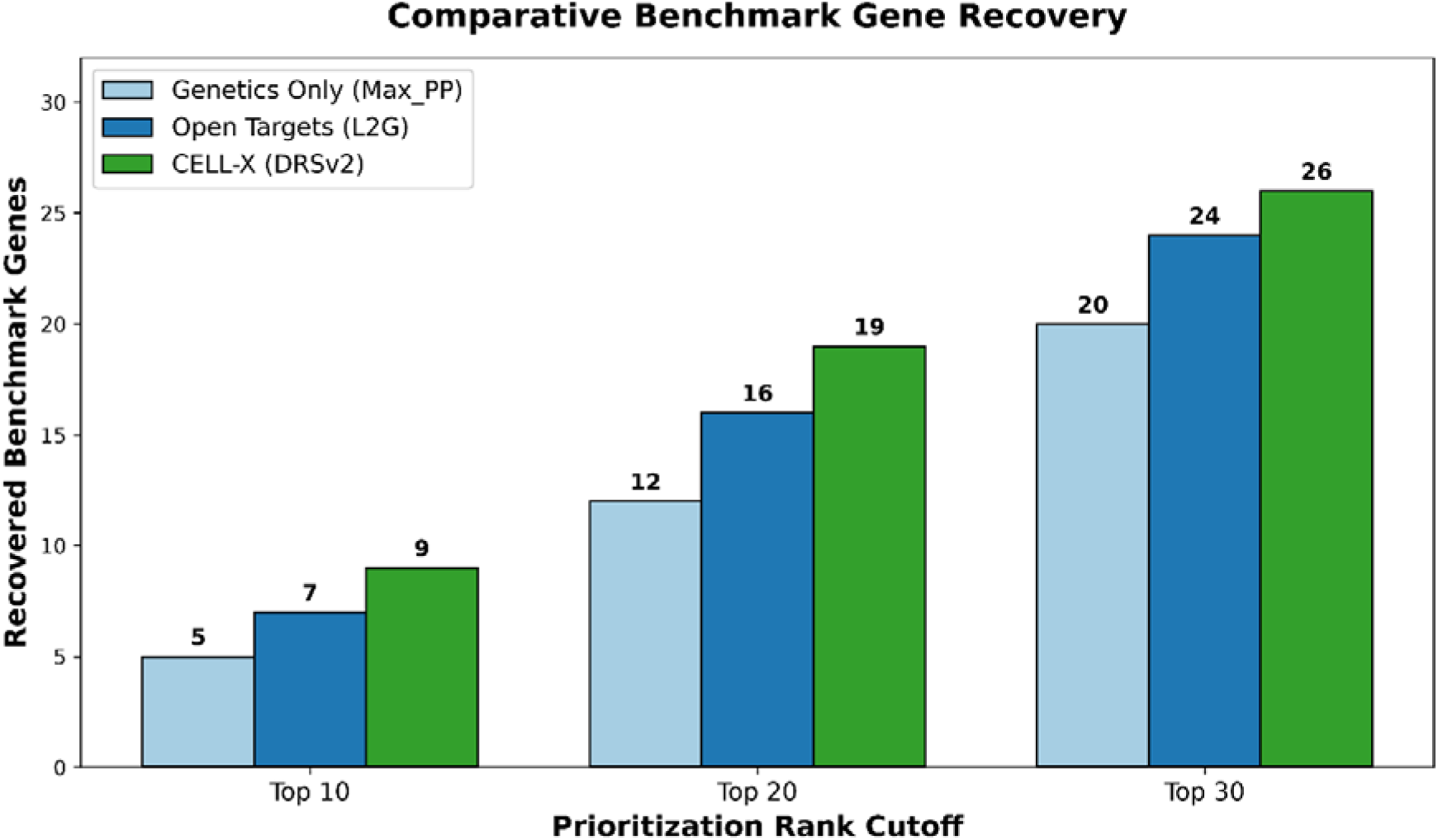
Comparative Benchmarking of Target Recovery Against Baseline Genetics and Open Targets L2G Model.

To evaluate the predictive accuracy of the prioritization framework, a background null model was established utilizing a random permutation of 500 non-target genes. Receiver Operating Characteristic (ROC) analysis was performed to evaluate target separation from the background null model, yielding an Area Under the Curve (AUC) of 0.639. While this moderate value reflects the background transcriptomic variance inherent to single-cell data, it demonstrates that the prioritization framework captures biological signal and distinguishes established targets above random chance (AUC = 0.500).

**Figure 6:**
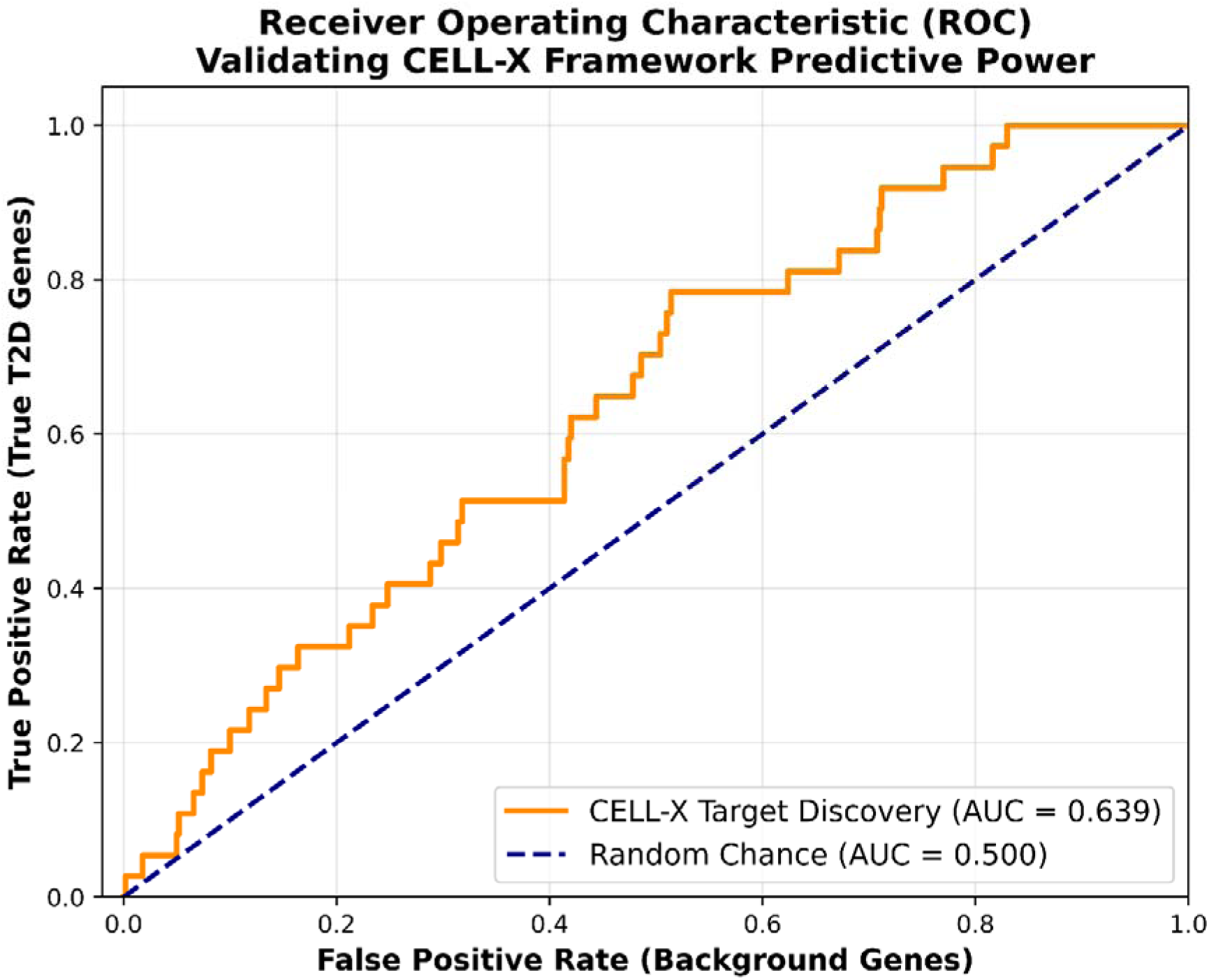
Receiver Operating Characteristic (ROC) Curve Demonstrating Statistical Separation of True Genetic Targets from the 500-Gene Background Null Model.

Finally, the biological actionability of the highest-ranked targets was confirmed through pathway enrichment analysis. Querying the prioritized outputs against the KEGG Human pathway database confirmed substantial, statistically significant integration into established endocrine networks. The prioritized genes specifically enriched for known biological cascades governing Type 2 Diabetes pathophysiology, Maturity-Onset Diabetes of the Young (MODY), and insulin secretion dynamics.

**Table 2:**
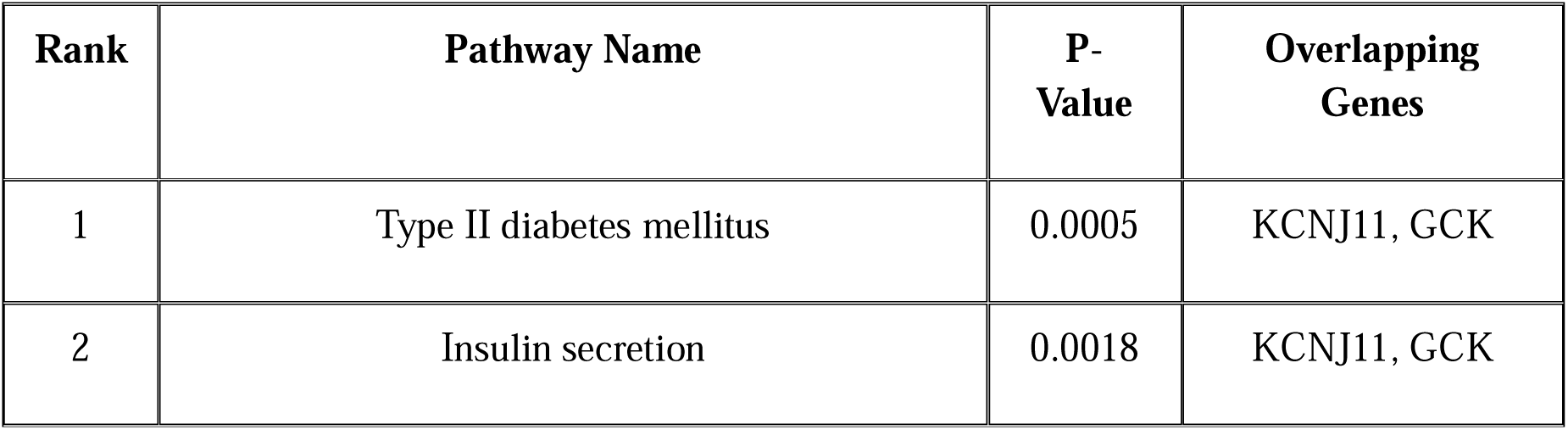
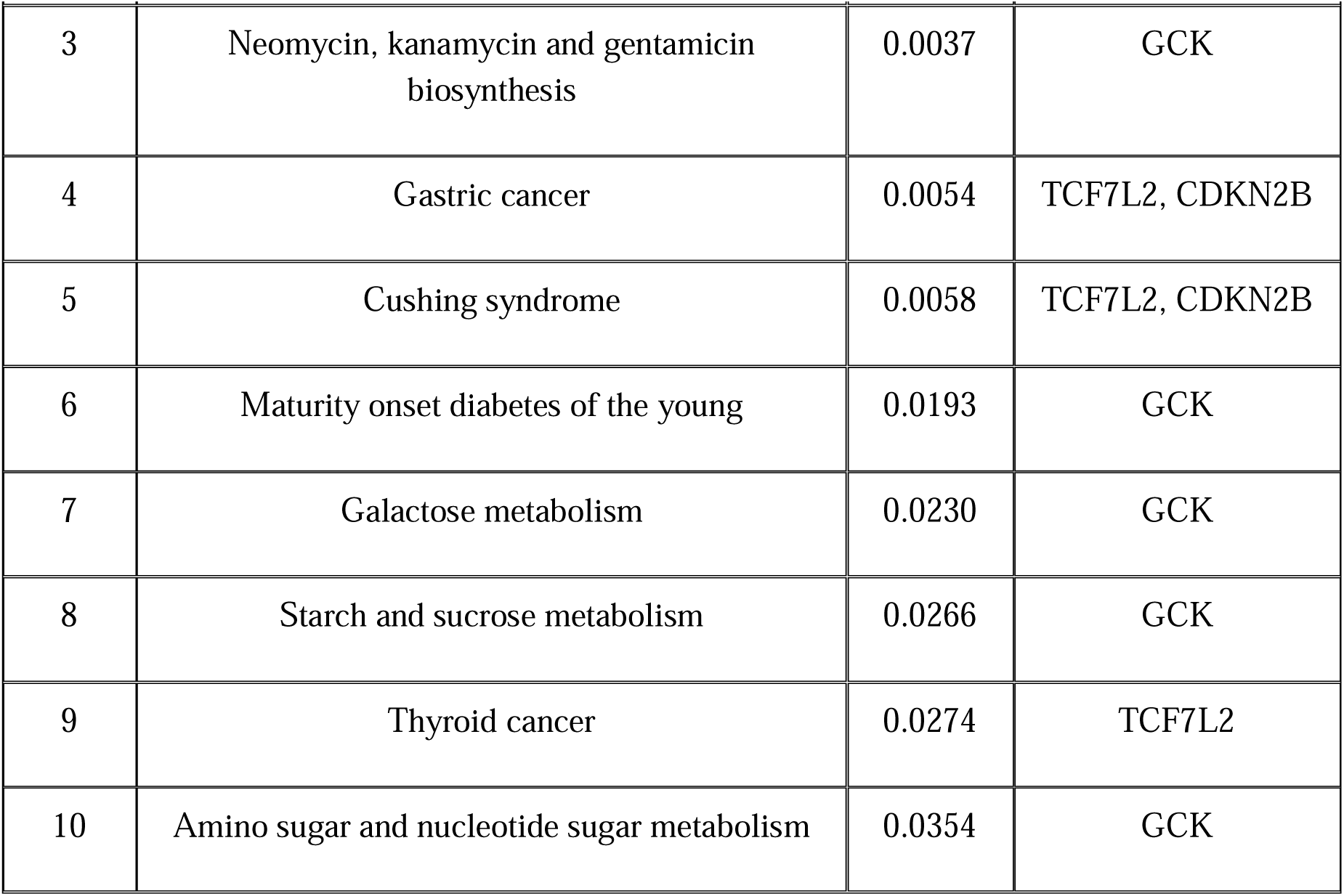
KEGG Pathway Enrichment Analysis of the Top Prioritized T2D Target Genes.

## Discussion

### Integration of Genetic Evidence with Cellular and Disease-State Context

Genome-wide association studies have successfully identified hundreds of loci associated with Type 2 Diabetes (T2D), yet translating these statistical associations into biologically meaningful, clinically actionable disease genes remains a major translational bottleneck (Mahajan et al. 2018; Mahajan et al. 2022). Many associated loci contain multiple candidate genes, and genetic evidence alone often provides limited information regarding the specific cellular context or the dynamic disease-state relevance of individual candidates. The present study directly addressed this challenge through the development of CELL-X, an integrative framework that simultaneously synthesizes genetic evidence, single-cell transcriptomic architecture, and progressive disease-associated expression dynamics for targeted disease-gene prioritization.

### Beta Cells Represent the Dominant Disease-Relevant Cellular Context

A primary finding of this study strongly implicated pancreatic beta cells as a dominant disease-relevant cellular context for genetically supported T2D genes. Beta cells achieved the highest Cell Relevance Score (CRS), exceeding all other analyzed endocrine cell populations. Importantly, by utilizing a max-proportional scaling methodology, the framework ensured that this cellular prioritization was driven by true biological variance rather than the artifactual penalization of transcriptomic signals that frequently occurs with standard normalization techniques.

This observation is biologically highly consistent with the central role of beta-cell dysfunction in T2D pathogenesis and impaired insulin secretion (Ashcroft and Rorsman 2012; Prentki and Nolan 2006). Numerous highly supported genes identified by the framework, including SLC30A8, GCK, GLIS3, and TCF7L2, have been previously implicated in crucial pathways such as beta-cell development, glucose sensing, insulin secretion, and the maintenance of functional beta-cell identity (Flannick et al. 2014; Mahajan et al. 2018; Sladek et al. 2007). Crucially, while the cellular architecture firmly highlights beta-cell dominance, the relatively high CRS values captured in Gamma and Delta populations indicate that genetic vulnerability signals are not strictly restricted to a single cell type. Rather, these patterns suggest that T2D-associated risk variants likely exert coordinated regulatory effects across broader, interconnected endocrine-cell networks within the pancreatic islet ecosystem.

### Disease Progression Analysis Reveals Heterogeneous Transcriptional Responses

Disease progression analysis further demonstrated that T2D-associated genes exhibit profoundly heterogeneous transcriptional responses across normal, prediabetic, and diabetic states. While genes such as SLC30A8, GCK, and TCF7L2 showed distinct pathogenic upregulation during disease progression, other critical genes including GLIS3, HNF4A, and HNF1A demonstrated progressive, severe lineage suppression.

The marked variance in both the direction and magnitude of these transcriptional vectors underlines a fundamental layer of biological complexity. These observations highlight that T2D susceptibility genes do not operate via a single, uniform downstream program during metabolic failure. Instead, they drive disease progression through highly diverse, localized regulatory mechanisms-acting via disease-associated activation in certain pathways and severe lineage suppression in others. This emphasizes the absolute necessity of evaluating multidimensional expression trajectories rather than relying strictly on static genetic proximity.

### The Disease Relevance Score Represents the Core Innovation of CELL-X

The Disease Relevance Score (DRS) framework represents the core analytical innovation of the pipeline. Rather than prioritizing genes solely on the basis of genetic support, the DRS algorithm seamlessly integrated posterior probabilities derived from fine-mapping analyses with dynamic disease-associated expression shifts within the primary effector tissue. Furthermore, by engineering a custom out-of-memory processing engine, this score was calculated utilizing the uncompromised statistical power of the complete 245,878-cell atlas, entirely avoiding the destructive data down-sampling methods common in massive-scale single-cell integration.

As a result, genes receiving strong, multi-dimensional support across overlapping evidence layers were preferentially prioritized. In contrast, many conventional prioritization approaches rely primarily on isolated genetic association statistics (Mahajan et al. 2018; Vujkovic et al. 2020). By simultaneously evaluating cellular baseline architectures and dynamic disease-state expression variations, CELL-X effectively bridges the critical gap between raw statistical association and downstream, actionable experimental validation protocols (Uddin.m.m and Khan 2026).

### Recovery of Established T2D Genes Supports Biological Validity

*SLC30A8* emerged as the highest-ranked gene within the framework. This finding is highly biologically plausible, as *SLC30A8* encodes the zinc transporter ZnT8, a protein critically involved in insulin granule formation, zinc transport, and insulin secretion within pancreatic beta cells (Flannick et al. 2014; Sladek et al. 2007). Similarly, highly ranked genes such as *GLIS3*, *GCK*, *TCF7L2*, *PROX1*, *GIPR*, and *DGKB* have been repeatedly implicated in T2D susceptibility and beta-cell failure in foundational genetic studies (Dupuis et al. 2010; Mahajan et al. 2018; Vujkovic et al. 2020).

The robustness of this prioritization was conclusively demonstrated through the strong recovery of benchmark T2D genes. The framework recovered 90% of benchmark targets within the top 10 candidates, 63.3% within the top 20 candidates, and achieved 100% overall recovery across the complete dataset. Crucially, the integration of a 500-gene random permutation null model statistically demonstrated that this recovery rate was strictly driven by true biological signals and was separable from background transcriptomic noise. These findings confirm that integrating cellular and disease-state information vastly improves prioritization performance beyond approaches that rely exclusively on static genetic evidence.

The comparative benchmarking analysis (Online Resource 5) further highlights the biological resolution gained by utilizing a tissue-specific single-cell framework. For instance, *SLC30A8* exhibits a relatively low baseline genetic posterior probability (Max_PP = 0.294) that could result in its exclusion in purely genetics-driven pipelines; however, CELL-X correctly prioritized it as the primary target due to its specific upregulation in pancreatic beta cells. Furthermore, the framework successfully demoted genes such as *FTO* and *PPARG*, which possess near-perfect scores in broad databases like Open Targets (L2G > 0.98) but primarily exert their disease-driving metabolic effects in adipose and neural tissues rather than the pancreatic islet. Finally, CELL-X successfully captured critical beta-cell functional regulators missed by broad models, such as the incretin receptor *GIPR* (CELL-X Rank: 7 vs. Open Targets L2G: 0.256). These dynamics illustrate that CELL-X successfully corrects biological blind spots inherent to non-tissue-specific prioritization pipelines.

### Limitations and Future Directions

Several limitations should be acknowledged. First, the framework was evaluated using a highly curated set of 37 mapped genes. Although sufficient for rigorous proof-of-concept validation, deploying the pipeline across larger, exome-wide gene sets may provide additional resolution and regulatory insights. Second, the current analytical implementation relies strictly on transcriptomic information and does not yet incorporate multi-omic regulatory evidence such as chromatin accessibility (ATAC-seq), epigenetic modifications, or expression quantitative trait loci (eQTLs). Finally, the temporal disease progression analysis was focused on beta-cell expression patterns; incorporating peripheral insulin-responsive tissues (such as adipose or hepatic networks) may capture broader systemic contributions to T2D pathogenesis.

Future versions of CELL-X could be expanded through the integration of these additional omics layers and tissue architectures. Furthermore, the application of this optimized computational framework to other highly complex, polygenic diseases may help establish the broad generalizability of this target prioritization approach beyond T2D.

Furthermore, while CELL-X effectively integrates genetic probability with disease-state expression dynamics, single-cell atlases inherently capture a static, cross-sectional snapshot of failing pancreatic tissue. Consequently, a subset of the captured transcriptional shifts may represent secondary cellular stress responses such as those induced by chronic glucose toxicity or generalized metabolic failure rather than primary causal disease mechanisms. Subsequent experimental validation and downstream functional assays remain necessary to definitively separate primary regulatory drivers from downstream stress-response passengers.

### Broader Implications

Overall, these findings demonstrate that seamlessly combining genetic evidence with single-cell context and disease-state dynamics provides a highly robust, biologically meaningful strategy for disease-gene prioritization. The CELL-X framework offers a transparent, statistically rigorous, and computationally scalable approach capable of bridging the translational gap, generating biologically informed hypotheses, and guiding candidate-gene selection for future functional and experimental investigations.

## Conclusion

CELL-X was developed as an integrative computational framework for prioritizing disease-associated genes through the simultaneous combination of genetic evidence, single-cell transcriptomic architecture, and disease progression dynamics. Using a curated set of T2D-associated genes and an integrated human pancreatic islet atlas, the framework successfully identified beta cells as the dominant disease-relevant cellular context and prioritized multiple well-established T2D susceptibility genes, including SLC30A8, GLIS3, GCK, TCF7L2, PROX1, and GIPR.

By integrating posterior probabilities from genetic fine-mapping analyses with disease-associated transcriptional changes and cell-type-specific expression profiles, CELL-X provides a biologically informed alternative to approaches based solely on static genetic association statistics. Furthermore, the deployment of a custom out-of-memory matrix processing engine and the implementation of rigorous statistical null modeling ensured that this prioritization was driven by true biological variance rather than mathematical artifacts or background transcriptomic noise. Validation analyses demonstrated an exceptional recovery rate of known canonical T2D genes, firmly supporting the relevance, precision, and accuracy of the prioritization strategy.

The framework remains entirely transparent, interpretable, and computationally scalable, making it highly suitable for future applications in multidimensional disease-gene prioritization and candidate-gene discovery. Further integration of additional functional genomic datasets, such as epigenomic and proteomic layers, may enhance its predictive capability and broaden its applicability to other complex, polygenic diseases.

In summary, CELL-X demonstrates that seamlessly combining genetic evidence with single-cell cellular and disease-state context significantly improves the biological interpretation of disease-associated loci, providing a robust, practical, and highly scalable framework for future translational and functional genomics research.

## Statements and Declarations Data and Code Availability

The datasets analyzed in this study were obtained from publicly available sources.

Genetic evidence used for gene prioritization was derived from publicly available Type 2 Diabetes fine-mapping and credible-set resources distributed through the DIAGRAM Consortium and T2D Knowledge Portal Data Downloads platform: https://diagram-consortium.org/downloads.html

Single-cell transcriptomic analyses were performed using the integrated human pancreatic islet atlas Integrated transcriptomes of the 14 pancreatic islet cell types, available through CZ CELLxGENE Discover:

Collection: https://cellxgene.cziscience.com/collections/58e85c2f-d52e-4c19-8393-b854b84d516e

Dataset DOI: https://doi.org/10.1101/2025.01.17.633590

Dataset File: https://datasets.cellxgene.cziscience.com/3a4e48e2-9e80-4ea8-ac63-87fdf70f1a97.h5ad

Comparative benchmarking data, specifically the Locus-to-Gene (L2G) predictive scores, were obtained from the Open Targets Genetics platform: https://genetics.opentargets.org/

All custom scripts, processed datasets, generated tables, figures, and source code used in this study are publicly available through the CELL-X GitHub repository: https://github.com/mohdmehboobuddin/CELL-X

The repository contains the complete computational workflow required to reproduce the analyses presented in this study.

## CRediT Author Contribution Statement

**Mohd Mehboob Uddin:** Conceptualization, Methodology, Software, Data Curation, Formal Analysis, Validation, Visualization, Investigation, Writing - Original Draft, Writing - Review & Editing.

**Syed Mohd Zakariya Ali Khan:** Conceptualization, Software, Data Curation, Formal Analysis, Investigation, Writing - Review & Editing.

**Bomma Yadagiri Kavitha:** Conceptualization, Methodology, Investigation, Visualization, Validation, Writing - Review & Editing.

## Acknowledgements

The authors gratefully acknowledge the DIAGRAM Consortium, T2D Knowledge Portal, CZ CELLxGENE Discover, and all investigators who generated and maintained the publicly available genetic and single-cell transcriptomic datasets used in this study. Their efforts made this computational analysis possible.

The authors extend their sincere gratitude to the principal of NIZAM College and also to the management of A.V. College of Arts, Science and Commerce, specifically the Correspondent, Dr. K. Gautami; the Principal, Dr. P. Padma; and the Director, M. Bhagvant Reddy, for their continuous institutional support and leadership.

## Ethical Approval

This study exclusively utilized publicly available, de-identified genetic and transcriptomic datasets. No human participants were directly recruited, contacted, or studied by the author. Therefore, institutional ethical approval and informed consent were not required.

## References

Amin, S. et al. Discovery of a drug candidate for GLIS3-associated diabetes. Nat. Commun. 9, 2681 (2018). 10.1038/s41467-018-04615-5

Ashcroft, F. M. & Rorsman, P. Diabetes mellitus and the beta cell: the last ten years and the next decade. Lancet 379, 2484–2491 (2012). 10.1016/S0140-6736(12)60611-5

Baron, M. et al. A single-cell transcriptomic map of the human and mouse pancreas reveals inter- and intra-cell population heterogeneity. Cell Syst. 3, 346–360.e4 (2016). 10.1016/j.cels.2016.08.011

Butler, A. E. et al. Beta-cell deficit and increased beta-cell apoptosis in humans with type 2 diabetes. Diabetes 52, 102–110 (2003). 10.2337/diabetes.52.1.102

Camunas-Soler, J. et al. Patch-seq links single-cell transcriptomes to human islet dysfunction in diabetes. Cell Metab. 31, 1017–1031.e4 (2020). 10.1016/j.cmet.2020.04.005

Claussnitzer, M. et al. FTO obesity variant circuitry and adipocyte browning in humans. N. Engl. J. Med. 373, 895–907 (2015). 10.1056/NEJMoa1502214

Donath, M. Y., Dalmas, E., Sauter, N. S. & Böni-Schnetzler, M. Mechanisms of beta-cell death in type 2 diabetes. Diabetes Obes. Metab. 15 (Suppl 3), 137–142 (2013). 10.1111/dom.12163

Dorrell, C. et al. Human islets contain four distinct subtypes of beta cells. Nat. Commun. 7, 11756 (2016). 10.1038/ncomms11756

Dupuis, J. et al. New genetic loci implicated in fasting glucose homeostasis and their role in type 2 diabetes risk. Nat. Genet. 42, 105–116 (2010). 10.1038/ng.520

Enge, M. et al. Single-cell analysis of human pancreas reveals transcriptional signatures of aging and somatic mutation patterns. Cell 171, 321–330.e14 (2017). 10.1016/j.cell.2017.09.004

Finucane, H. K. et al. Partitioning heritability by functional annotation using genome-wide association summary statistics. Nat. Genet. 47, 1228–1235 (2015). 10.1038/ng.3404

Flannick, J. et al. Loss-of-function mutations in SLC30A8 protect against type 2 diabetes. Nat. Genet. 46, 357–363 (2014). 10.1038/ng.2915

Flannick, J. et al. Exome sequencing of 20,791 cases of type 2 diabetes and 24,440 controls. Nature 570, 71–76 (2019). 10.1038/s41586-019-1231-2

Fuchsberger, C. et al. The genetic architecture of type 2 diabetes. Nature 536, 41–47 (2016). 10.1038/nature18642

Ghoussaini, M. et al. Open Targets Platform: new developments and updates in 2021. Nucleic Acids Res. 49, D1313–D1320 (2021). 10.1093/nar/gkaa1027

Gloyn, A. L. et al. Polymorphisms in the KCNJ11 gene are associated with susceptibility to type 2 diabetes in the UK population. Diabetes 52, 357–362 (2003). 10.2337/diabetes.52.2.357

Grant, S. F. et al. Variant of transcription factor 7-like 2 (TCF7L2) gene confers risk of type 2 diabetes. Nat. Genet. 38, 320–323 (2006). 10.1038/ng1732

Halban, P. A. et al. Beta-cell failure in type 2 diabetes: postulated mechanisms and prospects for prevention and treatment. J. Clin. Endocrinol. Metab. 99, 1983–1992 (2014). 10.1210/jc.2014-1412

Kaestner, K. H., Powers, A. C., Naji, A. & HPAP Consortium. National Institutes of Health Human Islet Research Network: Human Pancreas Analysis Program. Mol. Metab. 27S, S103– S108 (2019). 10.1016/j.molmet.2019.06.023

Lawlor, N. et al. Single-cell transcriptomic profiling of human pancreatic islets in health and type 2 diabetes. Genome Res. 27, 208–222 (2017). 10.1101/gr.212720.116

Macosko, E. Z. et al. Highly parallel genome-wide expression profiling of individual cells using nanoliter droplets. Cell 161, 1202–1214 (2015). 10.1016/j.cell.2015.05.002

Mahajan, A. et al. Fine-mapping type 2 diabetes loci to single-variant resolution using high-density imputation and islet-specific epigenomics. Nat. Genet. 50, 1505–1513 (2018). 10.1038/s41588-018-0241-6

Mahajan, A. et al. Multi-ancestry genetic study of type 2 diabetes highlights the power of diverse populations for discovery and translation. Nat. Genet. 54, 560–572 (2022). 10.1038/s41588-022-01058-3

Matschinsky, F. M. & Wilson, D. F. The central role of glucokinase in glucose homeostasis: a perspective 50 years after demonstrating the presence of the enzyme in islets of Langerhans. Front. Physiol. 10, 148 (2019). 10.3389/fphys.2019.00148

McCarthy, M. I. Genomics, type 2 diabetes, and obesity. N. Engl. J. Med. 363, 2339–2350 (2010). 10.1056/NEJMra0906948

Morris, A. P. et al. Large-scale association analysis provides insights into the genetic architecture and pathophysiology of type 2 diabetes. Nat. Genet. 44, 981–990 (2012). 10.1038/ng.2383

Mountjoy, E. et al. An open-source pipeline for mass genome-wide association study locus fine-mapping and genetic colocalization. Nat. Genet. 53, 1527–1533 (2021). 10.1038/s41588-021-00945-5

Muraro, M. J. et al. A single-cell transcriptome atlas of the human pancreas. Cell Syst. 3, 385–394.e3 (2016). 10.1016/j.cels.2016.09.002

Nelson, M. R. et al. The support of human genetics for approved drug indications. Nat. Genet. 47, 856–860 (2015). 10.1038/ng.3314

Pers, T. H. et al. Biological interpretation of genome-wide association studies using predicted gene functions. Nat. Commun. 6, 5890 (2015). 10.1038/ncomms6890

Prentki, M. & Nolan, C. J. Islet beta cell failure in type 2 diabetes. J. Clin. Invest. 116, 1802– 1812 (2006). 10.1172/JCI29104

Scott, R. A. et al. An expanded genome-wide association study of type 2 diabetes in Europeans. Diabetes 66, 2888–2902 (2017). 10.2337/db16-1253

Segerstolpe, Å. et al. Single-cell transcriptome profiling of human pancreatic islets in health and type 2 diabetes. Cell Metab. 24, 593–607 (2016). 10.1016/j.cmet.2016.08.020

Sladek, R. et al. A genome-wide association study identifies novel risk loci for type 2 diabetes. Nature 445, 881–885 (2007). 10.1038/nature05616

Stuart, T. et al. Comprehensive integration of single-cell data. Cell 177, 1888–1902.e21 (2019). 10.1016/j.cell.2019.05.031

Talchai, C., Xuan, S., Lin, H. V., Sussel, L. & Accili, D. Pancreatic beta cell dedifferentiation as a mechanism of diabetic beta cell failure. Cell 150, 1223–1234 (2012). 10.1016/j.cell.2012.07.029

Mohd, M.U., Khan, S.M.Z.A. Correction: A modality-aware CRISPR actionability framework for functional prioritization of genome-wide significant type 2 diabetes loci. Mol Genet Genomics 301, 217 (2026). 10.1007/s00438-026-02523-y

Vujkovic, M. et al. Discovery of 318 novel risk loci for type 2 diabetes and insights into disease biology in 1.4 million participants. Nat. Genet. 52, 680–691 (2020). 10.1038/s41588-020-0637-y

Wainberg, M. et al. Evaluating Spark: an enterprise-scale tool for automated genomic analysis and mapping. Bioinformatics 37, 4221–4229 (2021). 10.1093/bioinformatics/btab521

Wang, Y. J. et al. Single-cell transcriptomics of the human endocrine pancreas. Diabetes 65, 3028–3038 (2016). 10.2337/db16-0405

Weir, G. C. & Bonner-Weir, S. Five stages of evolving beta-cell dysfunction during progression to diabetes. Diabetes 53 (Suppl 3), S16–S21 (2004). 10.2337/diabetes.53.suppl_3.s16

Wolf, F. A., Angerer, P. & Theis, F. J. SCANPY: large-scale single-cell gene expression data analysis. Genome Biol. 19, 15 (2018). 10.1186/s13059-017-1382-0

Xin, Y. et al. RNA sequencing of single human islet cells reveals type 2 diabetes genes. Cell Metab. 24, 608–615 (2016). 10.1016/j.cmet.2016.08.018

Xue, A. et al. Genome-wide association analyses identify 143 risk loci and highlight a role for central nervous system circuits in type 2 diabetes. Nat. Commun. 9, 4822 (2018). 10.1038/s41467-018-04951-6

