## Supplementary Information for "CELL-X: A Single-Cell Transcriptomics Framework for Cell-Type and Disease-State Prioritization of Type 2 Diabetes Risk Genes"

**Author’s:**

Mohd Mehboob Uddin${}^{1}{}$*, Syed Mohd Zakariya Ali Khan${}^{1}{}$, BommaYadagiri Kavitha ${}^{2}{}$

**Affiliation’s:**

${}^{1}{}$Department of Life Sciences, A.V. College of Arts, Science and Commerce, Osmania University, Hyderabad 500029, Telangana, India

${}^{2}{}$Department of Biotechnology, A.V. College of Arts, Science and Commerce, Osmania University, Hyderabad 500029, Telangana, India

**Corresponding Author***

Mohd Mehboob Uddin

Department of Life Sciences, A.V. College of Arts, Science and Commerce, Osmania University, Hyderabad 500029, Telangana, India

**Overview**

This Online Resource-Supplementary Information document provides the extended datasets, comprehensive ranking tables, and additional data visualizations that support the findings presented in the main manuscript. The data contained herein details the baseline genetic evidence, the intermediate cell-type relevance mapping, and the comprehensive temporal expression dynamics across the full 37-gene target set utilized by the CELL-X framework.

**Online Resources (Supplementary Tables)**

**Online Resource 1:** Initial Genetic Evidence Dataset and Fine-Mapped GWAS Posterior Probabilities.

This table presents the genetic evidence dataset comprising the 37 mapped Type 2 Diabetes (T2D)-associated genes included in the CELL-X analysis. For each gene, the corresponding lead single nucleotide polymorphism (Top_SNP), maximum posterior probability (Max_PP), association p-value, and NCBI Gene ID are provided. The Max_PP value, derived from fine-mapping analyses, represents the genetic evidence supporting each candidate gene and serves as a foundational input for subsequent cell-type relevance and disease-state prioritization analyses.

| **Gene** | **Top SNP** | **Max_PP** | **P-Value** | **NCBI Gene ID** |
| --- | --- | --- | --- | --- |
| MTNR1B | rs10830963 | 0.9983 | 2.91E-12 | 4544 |
| HNF4A | rs1800961 | 0.9740 | 7.14E-06 | 3172 |
| GIPR | rs2238689 | 0.8992 | 2.46E-10 | 2696 |
| GCK | rs878521 | 0.8643 | 1.18E-09 | 2645 |
| KCNQ1 | rs74046911 | 0.8468 | 5.89E-18 | 3784 |
| TCF7L2 | rs7903146 | 0.7765 | 5.84E-120 | 6934 |
| HNF1A | rs1169288 | 0.7545 | 9.35E-09 | 6927 |
| GLIS3 | chr9:4294707:I | 0.5628 | 6.48E-08 | 169792 |
| CILP2 | rs72999033 | 0.5472 | 5.29E-13 | 148113 |
| CDKN2B | rs10811661 | 0.5451 | 3.25E-43 | 1030 |
| ZBED3 | rs7732130 | 0.4611 | 6.41E-10 | 114882 |
| PROX1 | rs340874 | 0.3522 | 1.12E-07 | 5629 |
| THADA | rs77981966 | 0.3147 | 1.74E-11 | 63892 |
| HNF1B | rs4430796 | 0.3129 | 6.31E-12 | 6928 |
| TSPAN8 | rs7961581 | 0.2954 | 1.15E-06 | 7103 |
| SLC30A8 | rs13266634 | 0.2946 | 1.28E-18 | 118430 |
| CDC123 | rs11257658 | 0.2866 | 2.92E-10 | 9022 |
| JAZF1 | rs1513272 | 0.2826 | 3.89E-12 | 22090 |
| CDKAL1 | rs35261542 | 0.2480 | 9.64E-23 | 54093 |
| ADCY5 | rs11708067 | 0.2287 | 1.14E-13 | 111 |
| GRB14 | chr2:165501927:I | 0.2232 | 2.37E-14 | 2888 |
| GCKR | rs780094 | 0.2231 | 8.50E-10 | 2646 |
| DGKB | rs10276674 | 0.1860 | 1.99E-05 | 1608 |
| BCL11A | rs243020 | 0.1835 | 2.46E-06 | 53335 |
| KCNJ11 | rs5215 | 0.1514 | 1.48E-10 | 3767 |
| ADAMTS9 | rs17676309 | 0.0982 | 1.17E-07 | 56999 |
| PPARG | rs11712037 | 0.0894 | 4.56E-08 | 5468 |
| IGF2BP2 | rs35510946 | 0.0884 | 3.70E-20 | 10644 |
| HHEX | rs11187140 | 0.0854 | 1.73E-14 | 3007 |
| IRS1 | rs2972156 | 0.0842 | 1.30E-15 | 3667 |
| PRC1 | rs3803563 | 0.0684 | 3.60E-10 | 9055 |
| WFS1 | rs10937721 | 0.0677 | 7.56E-11 | 7466 |
| HMGA2 | rs2583941 | 0.0668 | 7.49E-09 | 8091 |
| KLF14 | rs186463108 | 0.0514 | 4.55E-02 | 136259 |
| FTO | rs9927317 | 0.0435 | 1.66E-24 | 79068 |
| NOTCH2 | rs41463147 | 0.0257 | 5.83E-04 | 4853 |
| C2CD4B | rs138692931 | 0.0186 | 1.88E-03 | 388125 |

**Online Resource 2:** Complete Cell Relevance Scores (CRS) Across Major Pancreatic Endocrine Cell Populations.

This table provides the comprehensive output of the max-scaled Cell Relevance Score (CRS) calculations. The scores represent the relative spatial mapping of the genetically supported T2D gene set across the four primary endocrine cell populations: Beta, Gamma, Delta, and Alpha cells. By weighting the fine-mapped Max_PP values against the observed cell-type-specific expression fractions, this un-penalized, max-proportional scaling objective confirms that pancreatic beta cells exhibit the strongest cumulative transcriptomic enrichment for the prioritized targets, thereby justifying the framework’s subsequent focus on beta-cell disease-state dynamics.

| **Cell Type** | **Cell Relevance Score (CRS)** |
| --- | --- |
| **Beta** | 1.225 |
| **Gamma** | 1.142 |
| **Delta** | 1.104 |
| **Alpha** | 0.980 |

**Online Resource 3:** Temporal Disease Progression Expression Profiles of Prioritized Genes.

This dataset quantifies the raw transcriptomic shifts of the prioritized target genes across the metabolic disease continuum. The table lists the precise mean beta-cell expression values extracted from the integrated single-cell atlas, stratified across Normal (ND), Prediabetic (PD), and Type 2 Diabetes (T2D) clinical states. This temporal data highlights the underlying heterogeneity of T2D pathogenesis, revealing that targets experience distinct regulatory vectors (both pathogenic upregulation and severe lineage suppression) rather than a uniform metabolic response.

| **Gene** | **Normal (ND)** | **Prediabetes (PD)** | **Type 2 Diabetes (T2D)** |
| --- | --- | --- | --- |
| **MTNR1B** | 0.0000 | 0.0005 | 0.0007 |
| **HNF4A** | 0.0233 | 0.0153 | 0.0050 |
| **GIPR** | 0.0752 | 0.1227 | 0.0873 |
| **GCK** | 0.1656 | 0.1651 | 0.2299 |
| **KCNQ1** | 0.0026 | 0.0013 | 0.0012 |
| **TCF7L2** | 0.0779 | 0.0651 | 0.1140 |
| **HNF1A** | 0.0280 | 0.0202 | 0.0123 |
| **GLIS3** | 0.3479 | 0.2790 | 0.2712 |
| **CILP2** | 0.0001 | 0.0000 | 0.0000 |
| **CDKN2B** | 0.0277 | 0.0298 | 0.0565 |
| **ZBED3** | 0.0622 | 0.0858 | 0.0735 |
| **PROX1** | 0.1148 | 0.1287 | 0.1623 |
| **THADA** | 0.0443 | 0.0518 | 0.0496 |
| **HNF1B** | 0.0138 | 0.0113 | 0.0090 |
| **TSPAN8** | 0.0037 | 0.0167 | 0.0077 |
| **SLC30A8** | 1.2038 | 1.6101 | 1.4414 |
| **CDC123** | 0.2155 | 0.1943 | 0.2201 |
| **JAZF1** | 0.1543 | 0.1474 | 0.1794 |
| **CDKAL1** | 0.0393 | 0.0430 | 0.0468 |
| **ADCY5** | 0.0021 | 0.0062 | 0.0193 |
| **GRB14** | 0.0241 | 0.0125 | 0.0122 |
| **GCKR** | 0.0005 | 0.0002 | 0.0000 |
| **DGKB** | 0.0138 | 0.0462 | 0.1005 |
| **BCL11A** | 0.0125 | 0.0183 | 0.0077 |
| **KCNJ11** | 0.0784 | 0.0795 | 0.0880 |
| **ADAMTS9** | 0.0027 | 0.0012 | 0.0000 |
| **PPARG** | 0.0000 | 0.0007 | 0.0010 |
| **IGF2BP2** | 0.0197 | 0.0241 | 0.0308 |
| **HHEX** | 0.0053 | 0.0105 | 0.0150 |
| **IRS1** | 0.0072 | 0.0124 | 0.0131 |
| **PRC1** | 0.0052 | 0.0044 | 0.0055 |
| **WFS1** | 0.2441 | 0.2051 | 0.2243 |
| **HMGA2** | 0.0000 | 0.0002 | 0.0000 |
| **KLF14** | 0.0010 | 0.0007 | 0.0020 |
| **FTO** | 0.0615 | 0.0775 | 0.0844 |
| **NOTCH2** | 0.0043 | 0.0033 | 0.0010 |
| **C2CD4B** | 0.0458 | 0.2318 | 0.0455 |

**Online Resource 4:** Complete Disease Relevance Score (DRSv2) Rankings for the 37-Gene Target Set.

This table presents the final prioritization hierarchy for the entire curated gene set. The DRSv2 scores provided here represent the ultimate synthesis of the CELL-X framework, mathematically integrating the foundational genetic evidence (Max_PP), the absolute magnitude of the disease-state expression shift between T2D and ND states, and the overall targeted beta-cell expression. Genes are ranked in descending order of their DRSv2 scores, establishing a continuous spectrum of clinical actionability from highly supported canonical drivers (e.g., *SLC30A8*, *GLIS3*) down to transcriptionally inactive genetic neighbors.

| **Rank** | **Gene** | **Max_PP** | **Absolute Delta Shift (T2D vs ND)** | **Mean Beta Expression** | **DRSv2 Score** | **Direction of Effect** |
| --- | --- | --- | --- | --- | --- | --- |
| **1** | **SLC30A8** | **0.2946** | **0.2376** | **1.4184** | **9.93E-02** | **Pathogenic Upregulation** |
| **2** | **GLIS3** | **0.5628** | **0.0768** | **0.2994** | **1.29E-02** | **Lineage Suppression** |
| **3** | **GCK** | **0.8643** | **0.0643** | **0.1869** | **1.04E-02** | **Pathogenic Upregulation** |
| **4** | **TCF7L2** | **0.7765** | **0.0361** | **0.0857** | **2.40E-03** | **Pathogenic Upregulation** |
| **5** | **PROX1** | **0.3522** | **0.0475** | **0.1353** | **2.26E-03** | **Pathogenic Upregulation** |
| **6** | **JAZF1** | **0.2826** | **0.0251** | **0.1604** | **1.14E-03** | **Pathogenic Upregulation** |
| **7** | **GIPR** | **0.8992** | **0.0121** | **0.0950** | **1.03E-03** | **Pathogenic Upregulation** |
| **8** | **DGKB** | **0.1860** | **0.0868** | **0.0535** | **8.63E-04** | **Pathogenic Upregulation** |
| **9** | **CDKN2B** | **0.5451** | **0.0288** | **0.0380** | **5.97E-04** | **Pathogenic Upregulation** |
| **10** | **ZBED3** | **0.4611** | **0.0113** | **0.0738** | **3.86E-04** | **Pathogenic Upregulation** |
| **11** | **WFS1** | **0.0677** | **0.0198** | **0.2245** | **3.01E-04** | **Lineage Suppression** |
| **12** | **HNF4A** | **0.9740** | **0.0183** | **0.0145** | **2.59E-04** | **Lineage Suppression** |
| **13** | **CDC123** | **0.2866** | **0.0046** | **0.2100** | **2.74E-04** | **Pathogenic Upregulation** |
| **14** | **HNF1A** | **0.7545** | **0.0158** | **0.0202** | **2.40E-04** | **Lineage Suppression** |
| **15** | **KCNJ11** | **0.1514** | **0.0096** | **0.0820** | **1.19E-04** | **Pathogenic Upregulation** |
| **16** | **CDKAL1** | **0.2480** | **0.0075** | **0.0430** | **8.03E-05** | **Pathogenic Upregulation** |
| **17** | **THADA** | **0.3147** | **0.0052** | **0.0486** | **7.96E-05** | **Pathogenic Upregulation** |
| **18** | **FTO** | **0.0435** | **0.0230** | **0.0744** | **7.43E-05** | **Pathogenic Upregulation** |
| **19** | **GRB14** | **0.2232** | **0.0119** | **0.0163** | **4.31E-05** | **Lineage Suppression** |
| **20** | **ADCY5** | **0.2287** | **0.0171** | **0.0092** | **3.60E-05** | **Pathogenic Upregulation** |
| **21** | **IGF2BP2** | **0.0884** | **0.0112** | **0.0249** | **2.45E-05** | **Pathogenic Upregulation** |
| **22** | **HNF1B** | **0.3129** | **0.0048** | **0.0113** | **1.70E-05** | **Lineage Suppression** |
| **23** | **BCL11A** | **0.1835** | **0.0048** | **0.0129** | **1.14E-05** | **Lineage Suppression** |
| **24** | **TSPAN8** | **0.2954** | **0.0040** | **0.0094** | **1.10E-05** | **Pathogenic Upregulation** |
| **25** | **HHEX** | **0.0854** | **0.0097** | **0.0103** | **8.46E-06** | **Pathogenic Upregulation** |
| **26** | **IRS1** | **0.0842** | **0.0060** | **0.0109** | **5.51E-06** | **Pathogenic Upregulation** |
| **27** | **KCNQ1** | **0.8468** | **0.0014** | **0.0017** | **1.99E-06** | **Lineage Suppression** |
| **28** | **C2CD4B** | **0.0186** | **0.0003** | **0.1077** | **6.18E-07** | **Lineage Suppression** |
| **29** | **ADAMTS9** | **0.0982** | **0.0026** | **0.0013** | **3.33E-07** | **Lineage Suppression** |
| **30** | **MTNR1B** | **0.9983** | **0.0007** | **0.0004** | **2.81E-07** | **Pathogenic Upregulation** |
| **31** | **NOTCH2** | **0.0257** | **0.0033** | **0.0029** | **2.40E-07** | **Lineage Suppression** |
| **32** | **PRC1** | **0.0684** | **0.0003** | **0.0050** | **1.03E-07** | **Pathogenic Upregulation** |
| **33** | **KLF14** | **0.0514** | **0.0010** | **0.0012** | **6.55E-08** | **Pathogenic Upregulation** |
| **34** | **PPARG** | **0.0894** | **0.0010** | **0.0006** | **5.44E-08** | **Pathogenic Upregulation** |
| **35** | **GCKR** | **0.2231** | **0.0005** | **0.0002** | **2.80E-08** | **Lineage Suppression** |
| **36** | **CILP2** | **0.5472** | **0.0001** | **0.0000** | **1.98E-09** | **Lineage Suppression** |
| **37** | **HMGA2** | **0.0668** | **0.0000** | **0.0001** | **0.00E+00** | **Lineage Suppression** |

**Online Resource 5:** Comparative Benchmarking Scores against Baseline Genetics and Open Targets (L2G).

This table presents the comparative benchmarking data for the 37-gene CELL-X candidate set, incorporating baseline genetic evidence (Max_PP), Open Targets Genetics Locus-to-Gene (L2G) scores, and the final CELL-X Disease Relevance Score (DRSv2). The benchmark membership indicator identifies genes belonging to the predefined 30-gene Type 2 Diabetes susceptibility benchmark (1 = Benchmark Gene; 0 = Non-Benchmark Gene).

Across the prioritization thresholds evaluated, CELL-X recovered 9, 19, and 26 benchmark genes within the top 10, top 20, and top 30 candidates, respectively. The corresponding recovery counts were 5, 12, and 20 for baseline genetic prioritization using Max_PP, and 7, 16, and 24 for Open Targets L2G. These values provide the comparative recovery results presented in Figures 5A and 5B of the main manuscript.

| **Rank** | **Gene Symbol** | **Is Benchmark Gene** | **Genetics (Max_PP)** | **Open Targets (L2G)** | **CELL-X (DRSv2)** |
| --- | --- | --- | --- | --- | --- |
| **1** | **SLC30A8** | 1 | 0.2946 | 0.957 | 9.93E-02 |
| **2** | **GLIS3** | 1 | 0.5628 | 0.710 | 1.29E-02 |
| **3** | **GCK** | 1 | 0.8643 | 0.820 | 1.04E-02 |
| **4** | **TCF7L2** | 1 | 0.7765 | 0.885 | 2.40E-03 |
| **5** | **PROX1** | 1 | 0.3522 | 0.877 | 2.26E-03 |
| **6** | **JAZF1** | 1 | 0.2826 | 0.896 | 1.14E-03 |
| **7** | **GIPR** | 1 | 0.8992 | 0.256 | 1.03E-03 |
| **8** | **DGKB** | 1 | 0.1860 | 0.870 | 8.63E-04 |
| **9** | **CDKN2B** | 1 | 0.5451 | 0.572 | 5.97E-04 |
| **10** | **ZBED3** | 1 | 0.4611 | 0.582 | 3.86E-04 |
| **11** | **WFS1** | 1 | 0.0677 | 0.892 | 3.01E-04 |
| **12** | **CDC123** | 1 | 0.2866 | 0.743 | 2.74E-04 |
| **13** | **HNF4A** | 1 | 0.9740 | 0.901 | 2.59E-04 |
| **14** | **HNF1A** | 1 | 0.7545 | 0.967 | 2.40E-04 |
| **15** | **KCNJ11** | 1 | 0.1514 | 0.750 | 1.19E-04 |
| **16** | **CDKAL1** | 1 | 0.2480 | 0.834 | 8.03E-05 |
| **17** | **THADA** | 1 | 0.3147 | 0.815 | 7.96E-05 |
| **18** | **FTO** | 1 | 0.0435 | 0.984 | 7.43E-05 |
| **19** | **GRB14** | 1 | 0.2232 | 0.000 | 4.31E-05 |
| **20** | **ADCY5** | 1 | 0.2287 | 0.837 | 3.60E-05 |
| **21** | **IGF2BP2** | 1 | 0.0884 | 0.860 | 2.45E-05 |
| **22** | **HNF1B** | 1 | 0.3129 | 0.779 | 1.70E-05 |
| **23** | **BCL11A** | 1 | 0.1835 | 0.415 | 1.14E-05 |
| **24** | **TSPAN8** | 1 | 0.2954 | 0.683 | 1.10E-05 |
| **25** | **HHEX** | 0 | 0.0854 | 0.680 | 8.46E-06 |
| **26** | **IRS1** | 1 | 0.0842 | 0.654 | 5.51E-06 |
| **27** | **KCNQ1** | 1 | 0.8468 | 0.786 | 1.99E-06 |
| **28** | **C2CD4B** | 0 | 0.0186 | 0.349 | 6.18E-07 |
| **29** | **ADAMTS9** | 1 | 0.0982 | 0.812 | 3.33E-07 |
| **30** | **MTNR1B** | 0 | 0.9983 | 0.866 | 2.81E-07 |
| **31** | **NOTCH2** | 1 | 0.0257 | 0.729 | 2.40E-07 |
| **32** | **PRC1** | 0 | 0.0684 | 0.719 | 1.03E-07 |
| **33** | **KLF14** | 0 | 0.0514 | 0.090 | 6.55E-08 |
| **34** | **PPARG** | 1 | 0.0894 | 0.981 | 5.44E-08 |
| **35** | **GCKR** | 1 | 0.2231 | 0.785 | 2.80E-08 |
| **36** | **CILP2** | 0 | 0.5472 | 0.005 | 1.98E-09 |
| **37** | **HMGA2** | 0 | 0.0668 | 0.650 | 0.00E+00 |

**Online Resources (Supplementary Figures**)

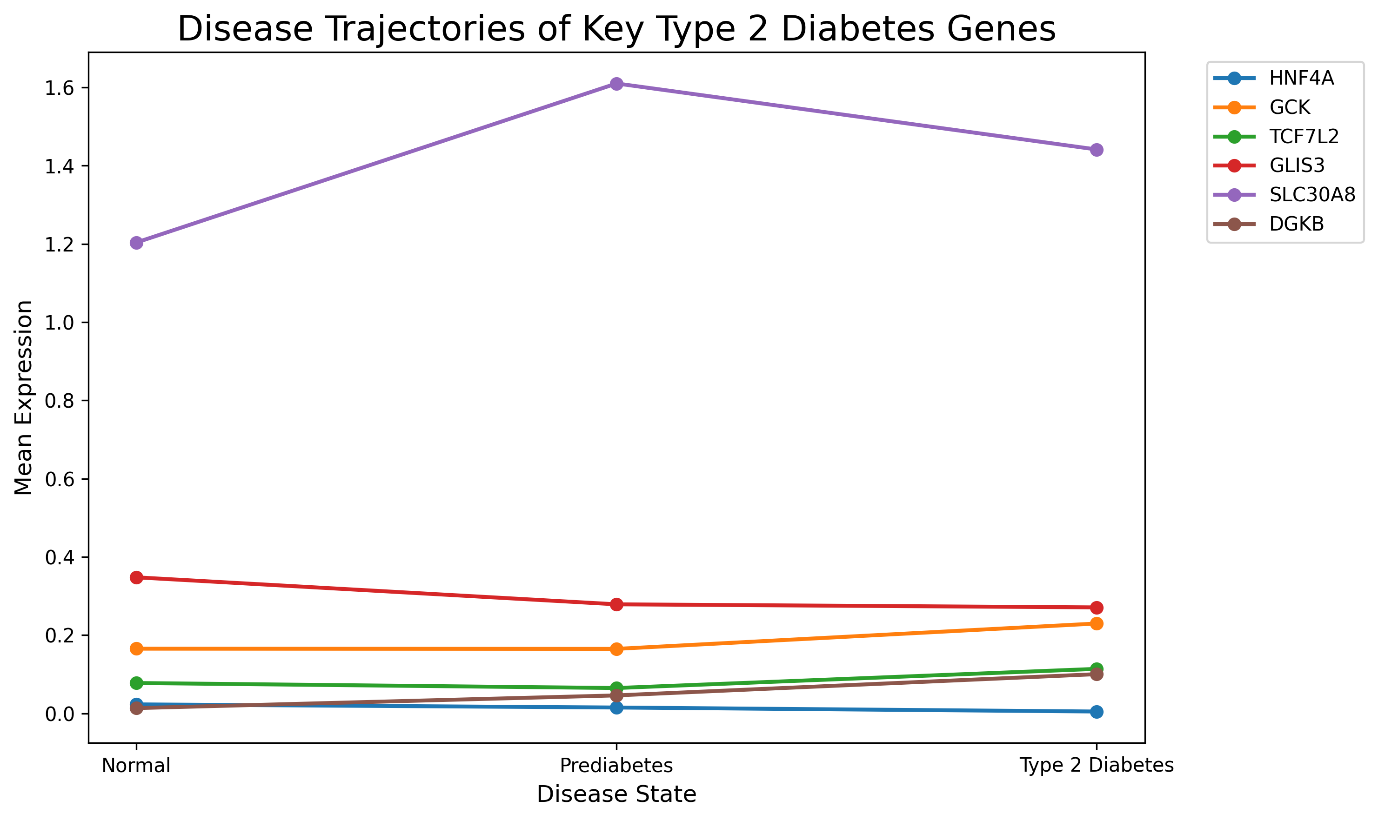

**Online Resource 6:** Gene-Specific Transcriptional Trajectories During Disease Progression.

Line plots illustrating the dynamic, multi-state temporal expression shifts of highly prioritized Type 2 Diabetes target genes. The x-axis represents the continuum of clinical metabolic states (Normal, Prediabetes, Type 2 Diabetes), while the y-axis denotes mean scaled expression within the beta-cell population. This visualization isolates the distinct pathogenic trajectories of key causal genes, highlighting continuous upregulation (e.g., *SLC30A8*) versus progressive lineage suppression (e.g., *GLIS3*, *HNF4A*).

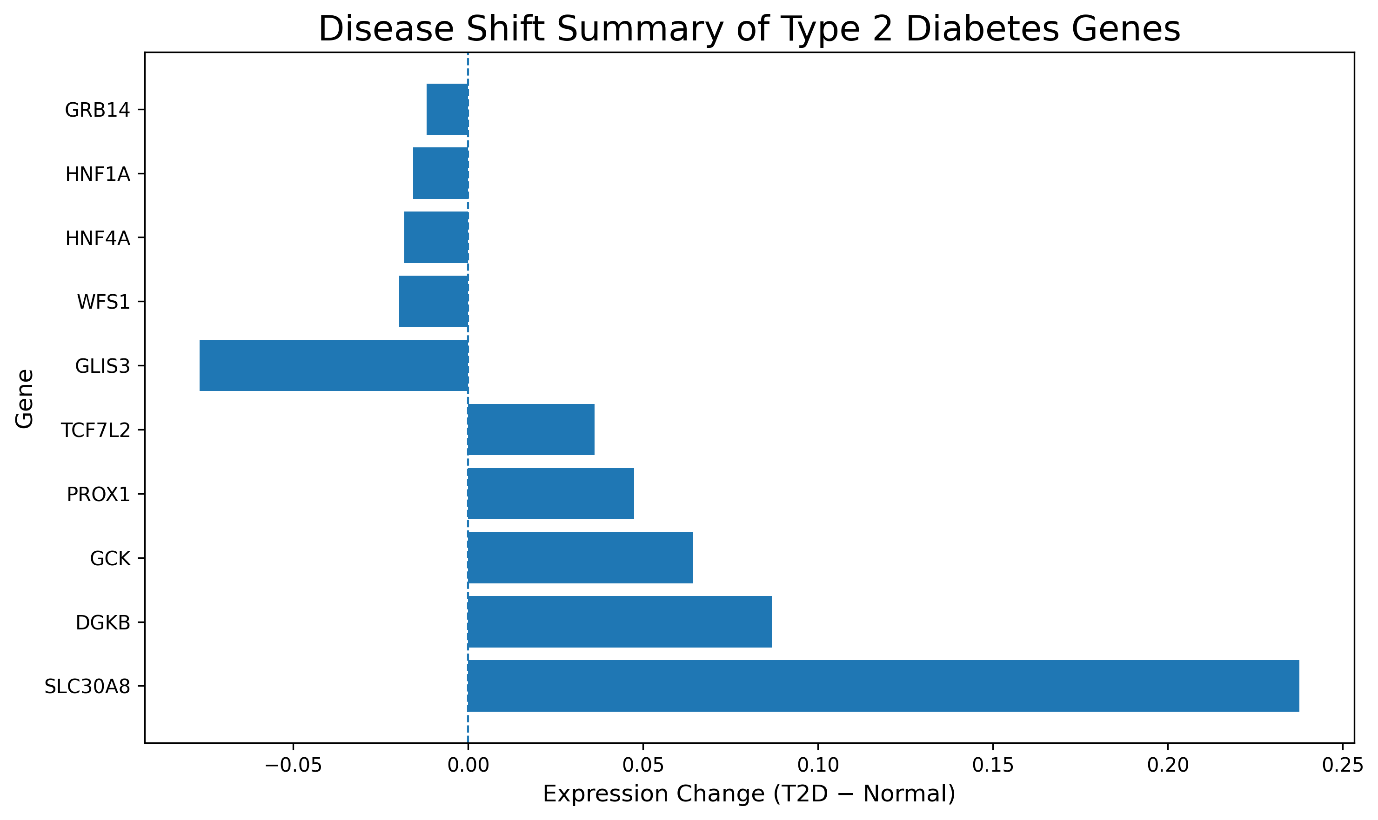

**Online Resource 7:** Absolute Magnitude and Directionality of Disease-Associated Expression Shifts.

Bar charts quantifying the absolute transcriptional change for the top prioritized genes. Values were calculated by subtracting the mean baseline beta-cell expression in Normal (ND) samples from the mean expression in Type 2 Diabetes (T2D) samples (Expression(T2D) - Expression(ND)). The distinct positive and negative vectors visually reinforce the highly heterogeneous regulatory responses occurring during metabolic failure.

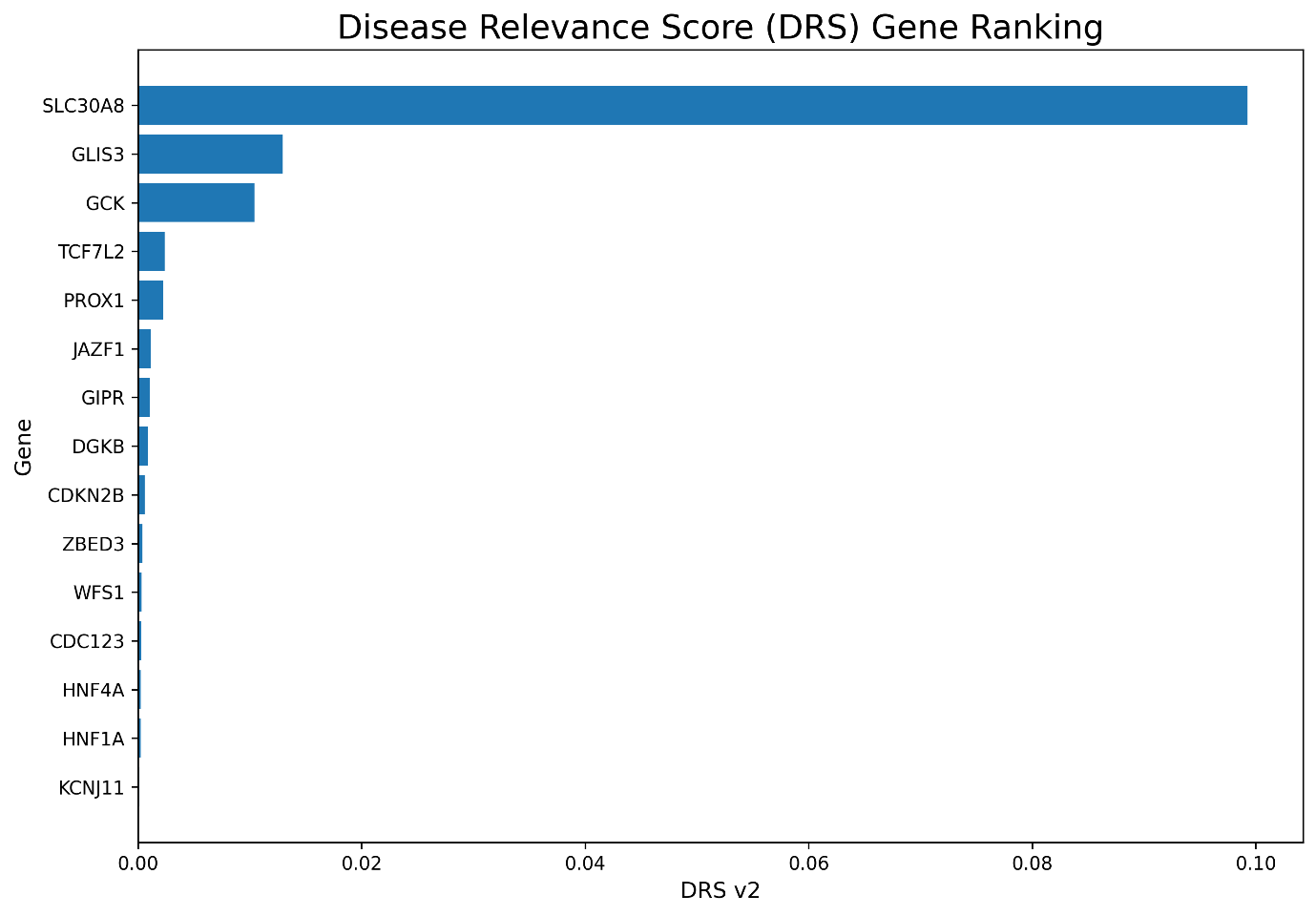

**Online Resource 8:** Top 15 Disease Relevance Score (DRSv2) Gene Prioritization.

This bar chart displays the 15 highest-ranked genes according to the final DRSv2 scores. It illustrates the separation between the most highly prioritized candidates and lower-ranked genes, while the complete 37-gene ranking is provided in Online Resource 4.

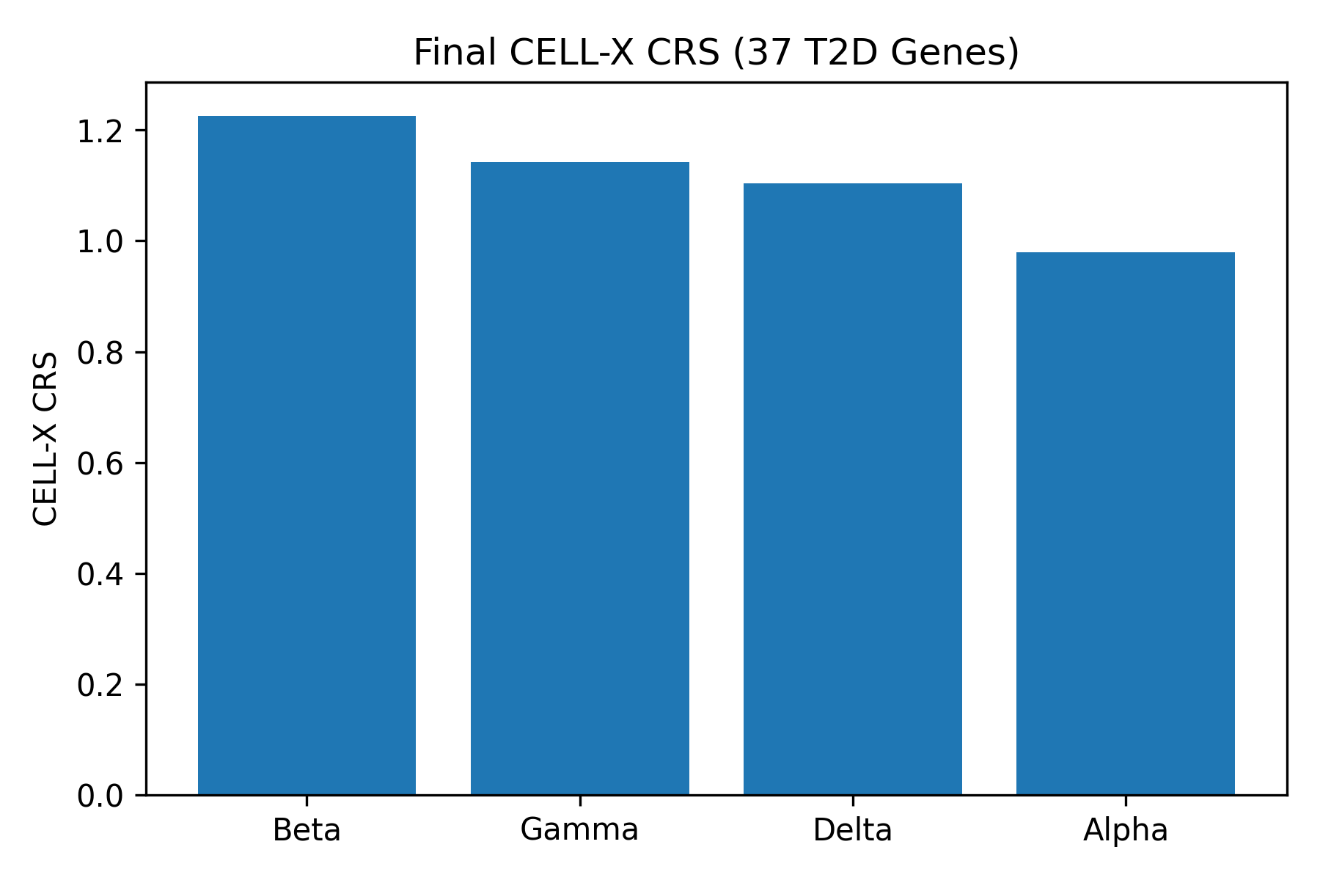

**Online Resource 9:** Final Spatial Relevance Mapping Across the Pancreatic Islet Atlas.

A detailed graphical breakdown of the cellular mapping and relevance scoring for the complete curated evidence dataset. The visualization confirms the robustness of the max-scaled Cell Relevance Score (CRS), visually reinforcing the beta-cell population as the dominant cellular context for the expression of T2D genetic risk variants.

**Online Resource 10:** Statistical Assessment of CELL-X Prioritization Using Background Null Modeling.

The statistical separation of prioritized Type 2 Diabetes candidate genes from background genes was evaluated using a random background set of 500 non-target genes processed through the CELL-X analytical framework. Receiver Operating Characteristic (ROC) analysis yielded an Area Under the Curve (AUC) of 0.639, compared with 0.500 for the random-classification reference. This result indicates modest separation between the evaluated target and background gene sets. The observed AUC provides an assessment of prioritization performance within the specified evaluation framework and does not, by itself, establish independent predictive validity.
